# Molecular Insights into the bactericidal Toxin Tle1 of *Pseudomonas aeruginosa*: Interaction with VgrG, its adaptor, and immunity protein

**DOI:** 10.64898/2025.12.24.696360

**Authors:** Delphine Lefebvre, Chantal Soscia, Laura Schmitt, Adeline Goulet, Bérengère Ize, Sophie Bleves

## Abstract

The Type VI secretion system (T6SS) delivers a wide range of antibacterial effectors, including phospholipases of the Tle family. Here, we characterize Tle1 from *Pseudomonas aeruginosa* and demonstrate that it functions as a bactericidal toxin with moderate activity when associated to the membranes in the periplasm. Bacterial two-hybrid assays revealed specific protein-protein interactions within the *tle1* locus, involving the immunity protein Tli1a, the chaperone/adaptor Tla1, and the spike protein VgrG4a. These interactions were independently validated by co-purification assays. Structural modeling with AlphaFold 3 produced a high-confidence ternary complex in which a VgrG4a trimer accommodates one Tle1 monomer and one Tla1 monomer. The three predicted interfaces (Tle1-Tla1, Tla1-VgrG4a, and Tle1-VgrG4a) were confirmed experimentally *in vivo* and important charged residues mediating these interfaces were identified. Furthermore, modeling of the Tle1-Tli1a complex suggests an inhibition mechanism that does not occlude the catalytic pocket. Consistently, Tli1a was localized to the outer membrane of *P. aeruginosa*, supporting *in silico* predictions of an outer membrane lipoprotein and positioning it ideally to neutralize periplasmic Tle1 activity. We did not observe the second candidate immunity protein, Tli1b, either in *P. aeruginosa* or in *E. coli*. Together, these findings elucidate the molecular interactions underlying Tle1 delivery and inhibition and highlight the role of Tli1a as a dedicated immunity protein that protects *P. aeruginosa* from self-intoxication.

## Introduction

*Pseudomonas aeruginosa* is one of the most feared opportunistic pathogens. This Gram-negative bacterium is associated with nosocomial infections, which are often severe and life-threatening, especially in immunocompromised hosts. Chronic infection leads to progressive lung disease in patients with cystic fibrosis. Antimicrobial resistance of *P. aeruginosa* is a predisposing factor for treatment failure and the biofilm mode of growth provides protection from antibiotics (1). The World Health Organization has classified *P. aeruginosa* in the high list of priority pathogens that pose the greatest threat to human health due to their resistance to antibiotics (2). *P. aeruginosa* has developed several pathogenicity strategies, of which protein secretion is a key one.

*P. aeruginosa* has three independent type VI secretion systems (T6SS) that allow it to compete with other bacteria or interact with eukaryotic hosts (3–5). The T6SS functions as a nanocrossbow that, upon contraction of a cytoplasmic sheath, releases a toxin-loaded arrow into the recipient cell where the toxins target conserved physiological processes. T6SS is a bacterial weapon capable of killing or inhibiting adjacent cells. The tube of the arrow is formed by a stack of Hcp (hemolysin co-regulated proteins), topped by VgrG (valine-glycine repeat protein G) and PAAR (proline-alanine-alanine-arginine) proteins, which form the spike complex (7). Two modes of secretion have been described so far. Firstly, T6SS toxins, also called effectors, can be either packaged inside the lumen of the Hcp tube or attached to the VgrG-PAAR spike, like a poisoned arrowhead. The spike-tip interaction of these cargo effectors can be direct or mediated by cytoplasmic chaperones also called adaptors, whose genes are encoded at the vicinity of effector genes (6). Secondly, evolved or specialized effectors, which are less frequently described, consist of a C-terminal effector extension of either Hcp, VgrG or PAAR proteins.

*P. aeruginosa* has a wide range of T6SS effectors, at least 22 in the PAO1 strain and 24 in the PA14 strain (4, 5, 8). In addition to secreting metallophores into the extracellular environment, *P. aeruginosa* injects toxins directly into competing bacteria, targeting various cellular components (peptidoglycan, cell membrane, DNA, proteins, and essential metabolic molecules). The gene encoding an antibacterial toxin is systematically close to a gene, or in some cases several copies, encoding an anti-toxin, also known as an immunity protein (9). The latter neutralizes the toxin activity in its compartment of action, protecting the bacterium from attack by sister cells or from self-intoxication in the case of toxins with activity in the cytoplasm.

Among these antibacterial toxins, one family is well represented in *P. aeruginosa*, the type VI lipase effectors (Tle) family, which hydrolyses membrane phospholipids (10). Tle toxins share very little sequence similarity and have been divided into five divergent families (Tle1-5) based on phylogeny and conserved catalytic motifs (10). Tle1 to 4 exhibit the GxSxG motif characteristic of lipases and esterases and are phospholipases A1 (PLA_1_) and/or PLA_2_, whereas members of the Tle5 family have a double HxKxxxxD motif common to phospholipases D. Outside of catalytic motifs, Tle toxins lack significant homology with known lipases. Type VI lipase immunity (Tli) genes encode proteins with lipoprotein or Sec signal sequence (SS), leading to the assumption that Tle effectors are active in the periplasm of the target bacterium.

The first Tle identified in *P. aeruginosa* is PldA (Tle5a), which is dependent on the H2-T6SS machinery (10). In the study that led to the discovery of PldB (Tle5b), these two effectors were shown to be both antibacterial and anti-eukaryotic, allowing *P. aeruginosa* to internalize into lung epithelial cells (11). These were the first examples of trans-kingdom toxins (5, 12). Interestingly three immunity proteins, presumably generated by gene duplication, have been described for PldB (Tle5b) and can individually protect against its toxicity (13). Subsequently, TplE (Tle4) was also described as a trans-kingdom PLA_1_ toxin, dependent on the H2-T6SS machinery, that triggers autophagy in infected cells and bacterial competition (14). Tle3 has then been described as antibacterial effector and requires a Tla3 chaperone to target it to VgrG2b of the H2-T6SS (15, 16). The 5^th^ Tle is Tle1, which has a PLA_2_ activity (17). Its structure has been solved by crystallography (17). Tle1 is organized into two distinct parts, the phospholipase catalytic domain (D1) with a α/β hydrolase fold and the membrane-anchoring module, composed of three amphipathic domains (D2, D3, D4). The same study revealed the toxicity of Tle1 produced in *Escherichia coli* depends on the Ser-Asp-His catalytic triad. However, its secretion mechanism in *P. aeruginosa* and its mode of protection/neutralization have never been characterized.

In this study, we demonstrate that Tle1 functions as a bactericidal toxin and provide molecular insights into its interaction network. Using AlphaFold 3-based structure prediction combined with experimental validation through two complementary protein-protein interaction assays, we decipher the specific interactions of Tle1 with its cognate immunity protein Tli1a, the adaptor protein Tla1, and the structural component of the T6SS machinery, VgrG4a.

## Results

### Tle1 is a bactericidal toxin

The antibacterial activity of Tle1 fused to a Sec SS in *E. coli* has been demonstrated by Hu and colleagues (17). We took advantage of this assay to determine whether Tle1 has bacteriostatic or bactericidal impacts on cell growth and viability. To answer this question, *tle1* sequence was cloned in frame with the sequence encoding the PelB SS and a C-terminal His-tag on the pET22b(+) vector under a P_T7_ promoter (called SS-Tle1_6His_, Table 1), and as a control, *tle1* was also cloned in the pET-Duet1 (called Tle1_6His_). The localization of these recombinant proteins in *E. coli* BL21(DE3) pLysS was verified by western-blot after cell fractionation (Fig. 1A). Tle1_6His_ produced in *E. coli* was recovered both in the cytoplasmic and the membrane fractions as the EF-Tu cytoplasmic protein and the Pal outer membrane lipoprotein controls, respectively (Fig. 1A, left panel). Interestingly SS-Tle1_6His_ was only found in the membrane fraction suggesting that when addressed to the periplasm via the SS, Tle1 targets the membrane (Fig. 1A, right panel). Figure 1B showed that whereas SS-Tle1_6His_ inhibits *E. coli* growth (4 log difference with control strain), Tle1_6His_ without a SS has no impact, demonstrating Tle1 toxicity from the periplasmic side of the membrane.

**Figure 1:**
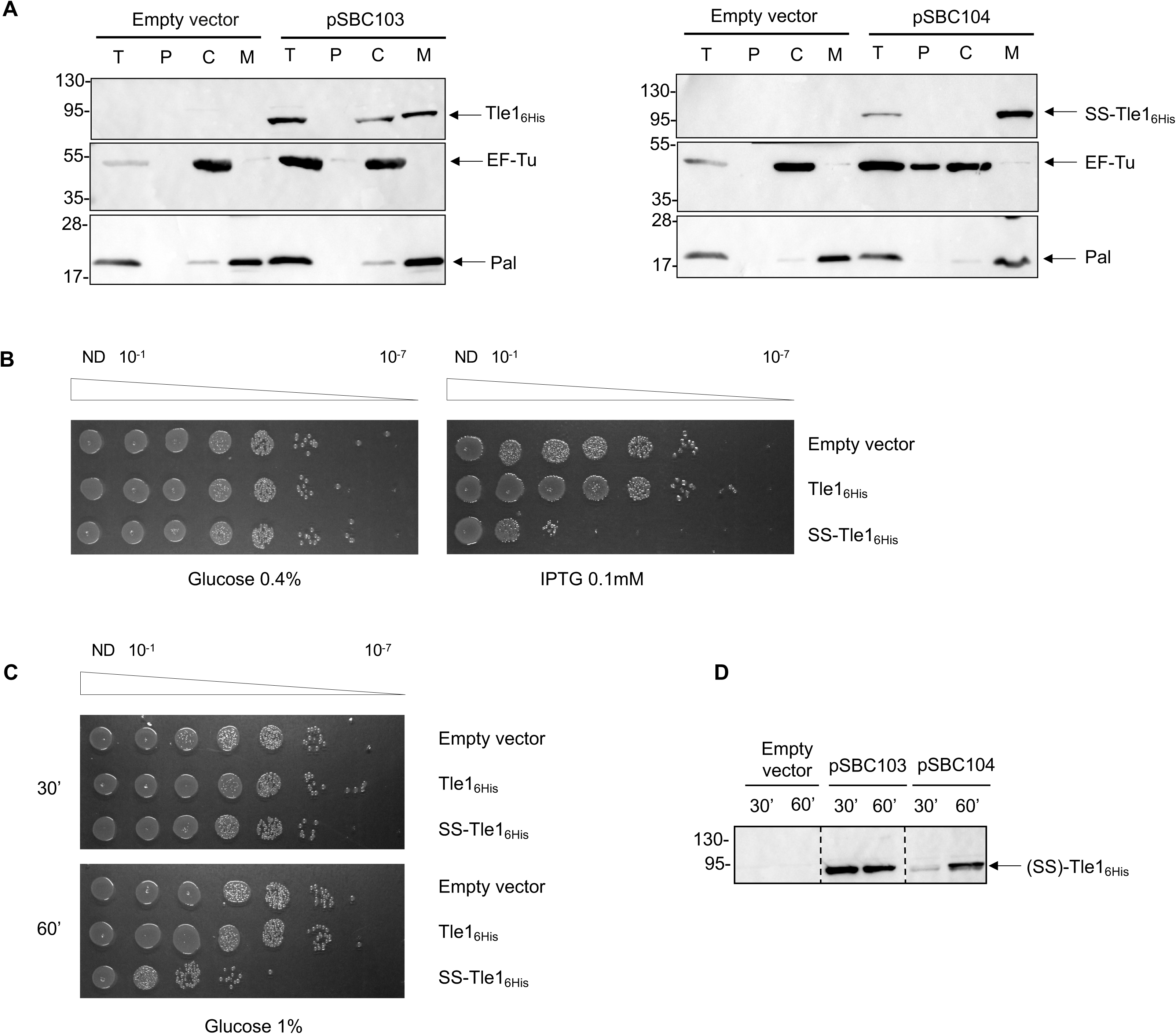
Tle1 is bactericidal towards *E. coli*. **(A) Tle1 fused to a Sec signal sequence is associated to membranes in *E. coli***. BL21(DE3) pLysS bacteria producing the wild-type Tle1, called Tle1_6His_ (from pSBC103, a pETDuet-1 derivative) or Tle1 fused to a Sec SS called SS-Tle1_6His_ (from pSBC104, a pET22b(+) derivative) were subjected to cell fractionation and immunoblotting T: bacteria, P: periplasm, M: membranes (IM and OM), C: cytoplasm. EF-Tu and Pal were used as cytoplasmic and membrane controls respectively. The position of the proteins and the molecular mass markers (in kDa) are indicated. It should be noted that production of SS-Tle1_6His_ promotes cell lysis. **(B) Tle1 is toxic when exposed to the periplasm**. Serial dilutions (from non-diluted (ND) to 10^−7^) of normalized cultures of *E. coli* BL21(DE3) pLysS producing Tle1_6His_, SS-Tle1_6His_, or carrying the empty vector were spotted on LB agar plates supplemented with 0.4% glucose (left panel) or with 0.1 mM IPTG (right panel). Glucose and IPTG allow respectively repression and induction of the gene encoding the T7 RNA polymerase. **(C) Bactericidal effect is associated with the production of SS-Tle1**. *E. coli* BL21(DE3) pLysS producing Tle1_6His_, SS-Tle1_6His_, or carrying the empty vector were harvested 30- and 60 -min post-induction by 0.1 mM IPTG. Serial dilutions of normalized cultures were spotted on LB agar containing 1% glucose to repress the production of the indicated protein. **(D) Immunodetection of Tle1 and SS-Tle1 from (C).** Cells from panel (C) were collected at each time point and analyzed by Western blot using an anti-His antibody.

**Table 1:**
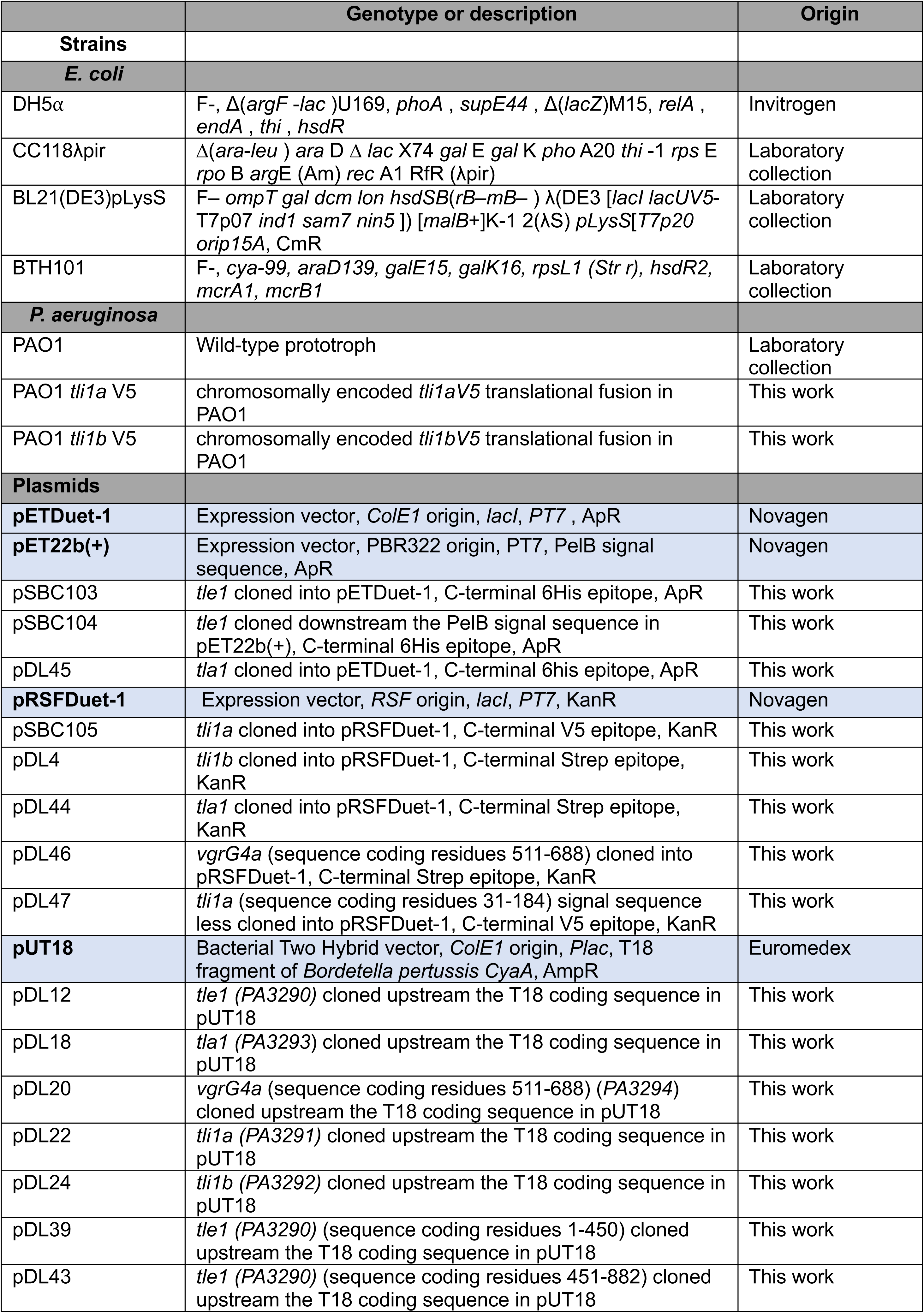
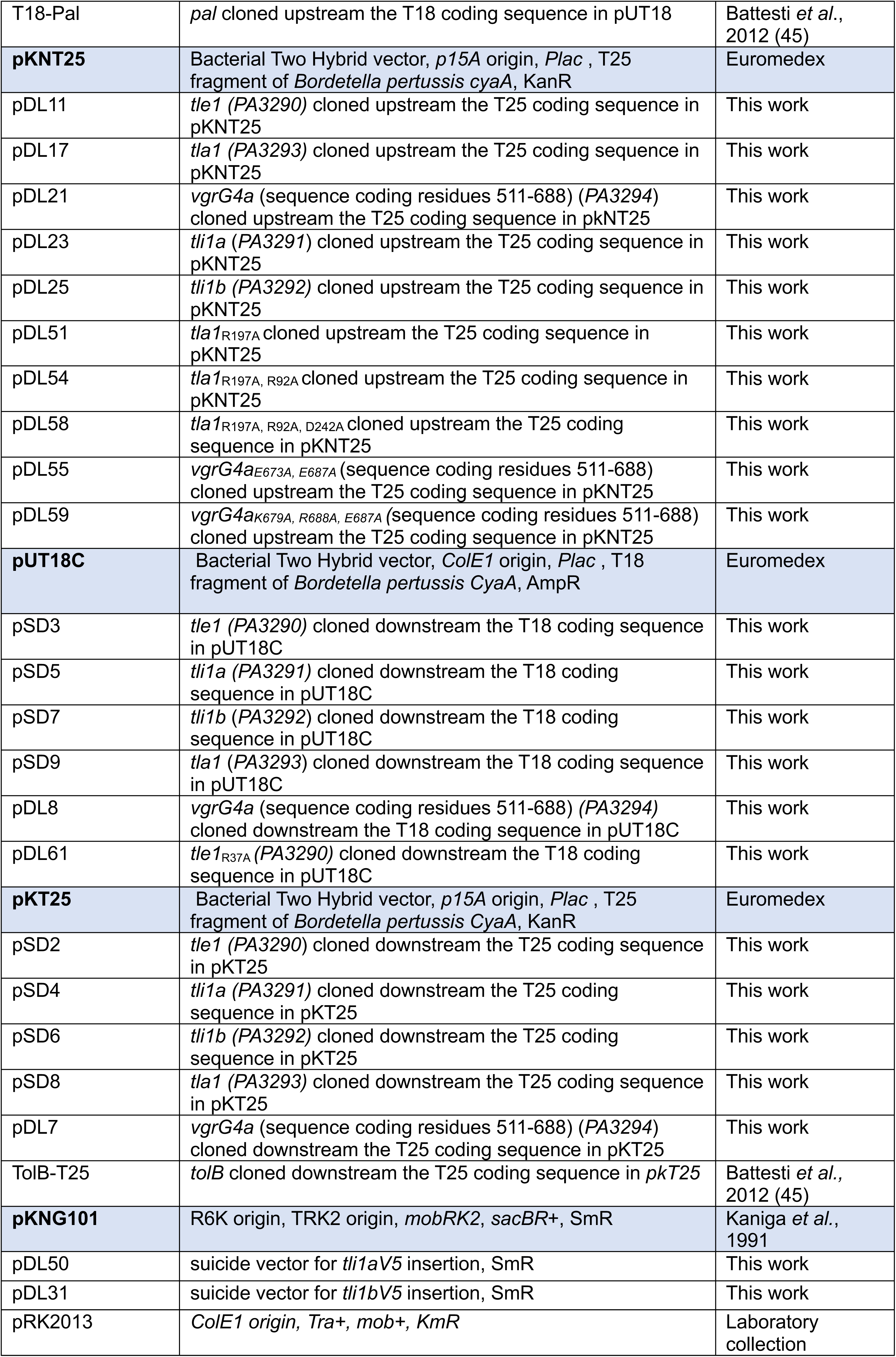
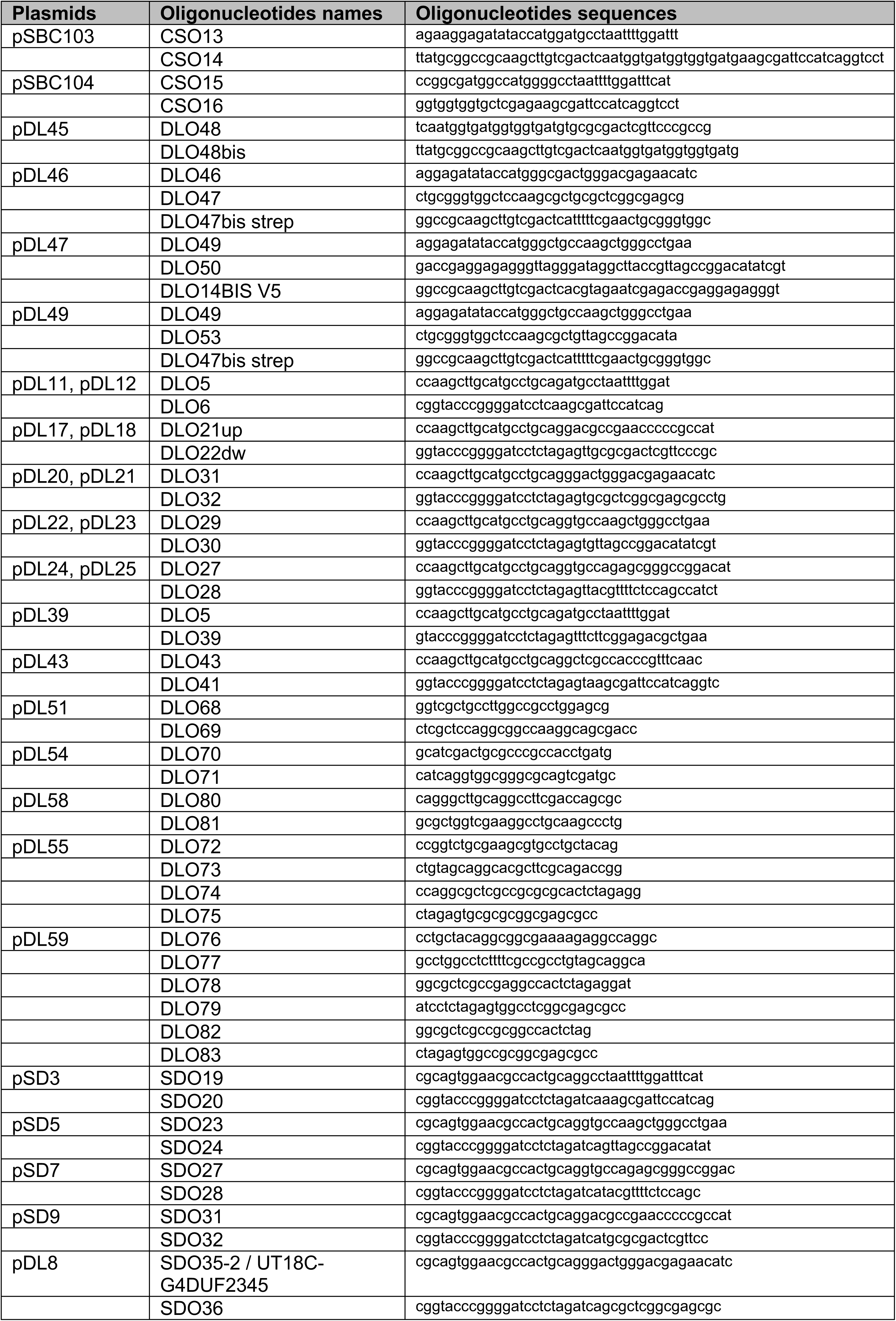
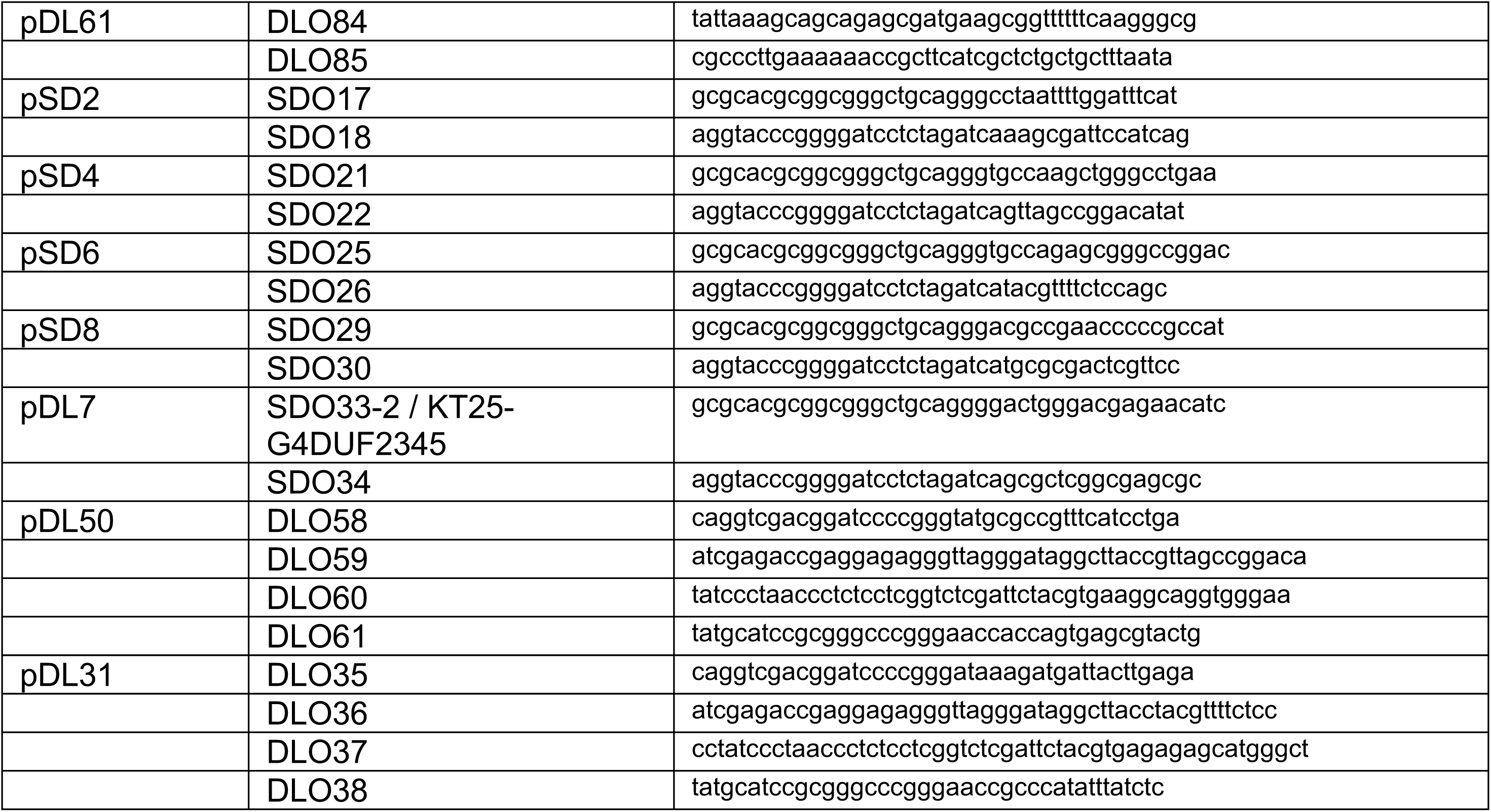
*E. coli* and *P. aeruginosa* strains, plasmids and oligonucleotides used in this study.

The bactericidal/bacteriostatic assay was then undertaken (18). The aim was to observe whether bacteria could resume growth (bacteriostatic effect) or not (bactericidal effect) after Tle1 toxic form was produced in *E. coli*. Bacteria were then grown in liquid medium, cells were harvested at 30- and 60-minutes post-induction of *ss-tle1_6His_* or *tle1_6His_*, washed and after normalization to OD_600_, spotted on a repressive LB agar medium (Fig. 1C). In figure 1C, bacteria at 30 min of induction of *ss-tle1_6His_* were able to grow on a repressive medium as Tle1_6His_ or the negative control strain. But the immunodetection of SS-Tle1_6His_ in these bacteria revealed it was slightly produced in comparison to Tle1_6His_ (Fig. 1D). After 60 min of SS-Tle1_6His_ production, bacteria were no more able to resume growth, and the toxin was clearly immunodetected. Taken together, these results indicate that Tle1 is a bactericidal toxin with moderate activity.

### Tle1 protein-protein interaction network

The analysis of Tle1 genetic environment in *P. aeruginosa* PAO1 strain revealed the presence of 4 other genes (Fig. 2A). PA3294 encodes VgrG4a, a protein belonging to the puncturing tip of a T6SS machinery. In *P. aeruginosa*, in addition to the 3 clusters of genes encoding the core components of the T6SS machineries, there are 8 so-called orphan islands containing *vgrG* and/or *hcp* genes scattered around the chromosome and encoding structural elements of the T6SS (19). They are considered predictive of T6SS effector candidates because genes encoding effectors and those involved in their secretion are associated with these loci. The *tle1* gene belongs to such an operon localized at distance from the T6SS core genes. Next to *vgrG4a*, PA3293 encodes a predicted cytoplasmic protein with a DUF4123 domain, found in T6SS chaperone/adaptor proteins responsible for addressing effectors to the machinery in the cytoplasm (3, 20) and has been named Tla1 for Type VI lipase adaptor protein. Effector genes are usually encoded with a cognate immunity gene avoiding a self-toxic activity and/or preventing T6SS-dependent killing by neighboring cells. The two next genes (PA3291 and PA3292) adjacent to the *tle1* gene are good candidates for encoding Tle1 immunity proteins and have therefore been named Tli1a and Tli1b respectively. SignalP 6.0 predictions indicate that they both harbor a lipoprotein signal sequence (Fig. S1). According to the two known lipoproteins sorting signals (the Lol avoidance signal (aspartate at position +2 of the mature protein) (21), and the +3, +4 rule of *P. aeruginosa* lipoproteins (22)), Tli1a and Tli1b would be anchored in the outer membrane. Consequently, both proteins may be exposed to the periplasm, which is consistent with the activity of Tle1 in this compartment (Fig. 1). Protein sequence analysis using BLASTP revealed that Tli1a and Tli1b are 75% identical over almost 92% of the total Tli1b sequence (Fig. S2A). Both proteins contain at least one domain of unknown function (DUF3304), which is found in immunity proteins neutralizing Tle1 effectors from three different bacteria: *Burkholderia thailandensis*, *Klebsiella pneumoniae,* and *Vibrio cholerae* (10, 23, 24). Tli1a contains a single DUF3304, whereas Tli1b has a full-length DUF3304 and a shorter one in its C-terminal (Fig. 2A). BLAST shows 75% identity with the full-length DUF3304 of Tli1b and 57% with the shorter DUF3304, suggesting a possible duplication (Fig. S2B).

**Figure 2:**
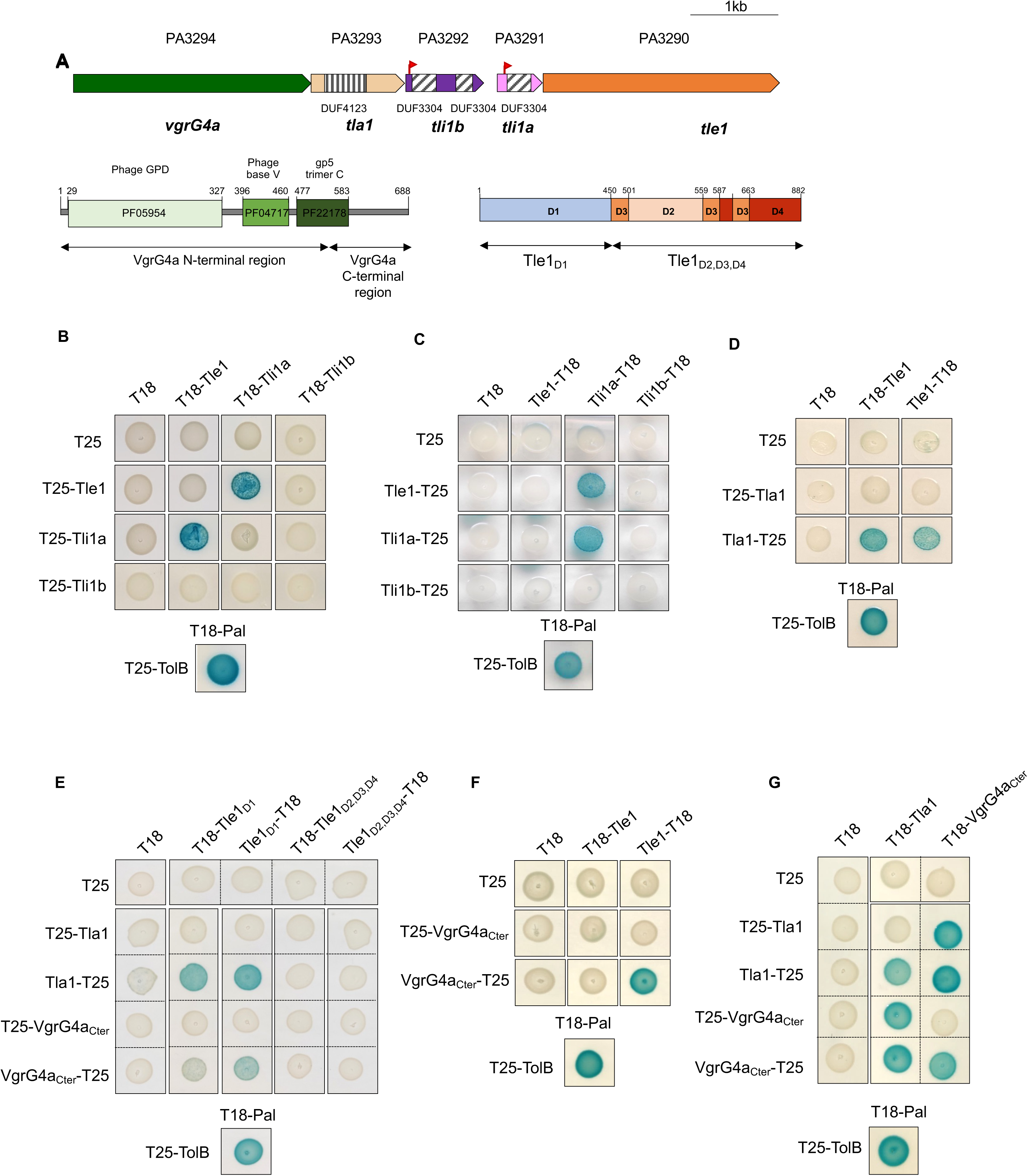
Tle1 interaction network. **(A) *vgrG4a* island organization.** The genes are labelled with the given name (i.e., *tle1*) and are indicated by their annotation number (e.g., PA3290). The lipoprotein signal peptides of Tli1a (PA3291) and Tli1b (PA3292) are represented with red flags, the DUF4123 of Tla1 (PA3293) with striped box (vertical) and DUF3304 of Tli1a and Tli1b with striped boxes (oblique). The domain architecture organization of VgrG4a and Tle1 into domains is presented under the cognate gene. **(B-G) Bacterial two-hybrid assay.** BTH101 reporter cells producing the indicated proteins or domains fused to the T18 or T25 domain of the *B. pertussis* adenylate cyclase were spotted on X-gal indicator plates. The blue colour of the colony reflects the interaction between the two proteins. TolB and Pal are two proteins known to interact but unrelated to the T6SS. The experiment was performed in triplicate and a representative result is shown.

To decipher the secretion mechanism of Tle1, we performed a bacterial two-hybrid (BACTH) assay with the other gene products of the locus, hypothesizing that a genetic link could reflect protein-protein interactions. The sequences encoding Tle1, Tla1, Tli1a and Tli1b excluding their SS and the C-terminal domain of VgrG4a (Fig. 2A) were cloned either downstream or upstream the sequences encoding T18 or T25 domains of the *Bordetella* adenylate cyclase. VgrG4a harbors a gp27-like hub domain, followed by a gp5 C-terminal domain and a C-terminal extension, an organization typical of VgrG proteins (Fig. 2A). Since we and others have delimited the interaction domain of two Tle to the C-terminal domain of VgrGs (15, 25), we cloned only this sequence of VgrG4a (VgrG4a_Cter_).

The BACTH assay revealed interactions between Tle1 and Tli1a (Fig. 2B and 2C) since coproduction of the T18/T25-Tle1 and T18/T25-Tli1a fusion proteins and Tle1-T18 and Tli1a-T25 combinations activated the expression of the *lacZ* reporter gene, as the positive control of the Tol-Pal interaction (26). At least dimerization of Tli1a was observed when the N-terminal domain of Tli1a was free (compare Fig. 2B and 2C). Unexpectedly, Tle1 did not interact in this assay with Tli1b in any of the orientations tested (Fig. 2B and 2C). Then, since Tle1 has two distinct domains, we wondered whether Tli1a binds preferentially to one of them (Fig. 2A). Interestingly, the use of either the catalytic domain (D1) or the membrane-anchoring module of Tle1 (D2, D3 and D4) disrupts the Tle1-Tli1a interaction in BACTH (Fig. S3). This suggests that both domains of Tle1 are important for Tli1a binding to Tle1.

The Figure 2D showed an interaction between Tle1 and Tla1. Interestingly, the T25-Tla1 fusion failed to interact with T18-Tle1 or Tle1-T18, while this fusion is active for an interaction with VgrG4a (Fig. 2G). This suggests that the fusion of the T25 domain to the N-terminal of Tla1 affects the Tle1-Tla1 interaction. It is thus possible that fusing the T25 domain to the N-terminus of Tla1 (the two proteins are almost the same size) masks the interaction interface with Tle1, while it would not be the case for fusion at the C-terminus. The interaction domain of Tle1 with Tla1 was further delimitated to its catalytic domain (Fig. 2E). Furthermore, Tle1 was also able to interact with the T6SS puncturing protein VgrG4a (Fig. 2F). This interaction requires the N-terminal catalytic domain of Tle1 (Fig. 2E).

Finally, to confirm the functions of Tla1 and VgrG4a_Cter_, their interaction was tested (Fig. 2G). Both proteins interacted with each other consistent with an adaptor protein function for Tla1 that interacts with a VgrG protein of the perforation tip to address its cognate effector. The two proteins can at least dimerize (Fig. 2G) which agrees with the VgrG trimers described in the literature (27, 28).

We were thus able to detect, by BACTH, interactions between proteins of the *tle1* locus, except for Tli1b. These results are consistent with our hypotheses of an adaptor protein function for Tla1 that interacts with both the effector Tle1 and the structural component VgrG4a, and an immunity protein function for Tli1a.

### How many immunities for Tle1?

To confirm the interaction between Tle1 and Tli1a, we undertook a complementary co-purification approach of affinity chromatography. A cytoplasmic V5-tagged version of Tli1a was engineered by fusing the tag to the mature domain of Tli1a, lacking its lipoprotein SS. The recombinant protein was co-produced in *E. coli* BL21(DE3) pLysS with Tle1_6His_, the not toxic form of Tle1 from the bactericidal assay (Fig. 1), and an affinity chromatography on nickel matrix was performed (Fig. 3A). The presence of Tli1a_V5_ was controlled in the elution fraction with V5 antibodies. As showed in figure 3A, Tli1a_V5_ was found in the eluted fraction only upon coproduction with Tle1_6His_ (left panel). Indeed, when produced alone in *E. coli*, Tli1a_V5_ was not purified by affinity chromatography (right panel). As expected for an immunity protein, Tli1a directly interacts with Tle1. A V5-tagged cytoplasmic form of Tli1b was also generated. However, this recombinant form was not detectable in western blots.

**Figure 3:**
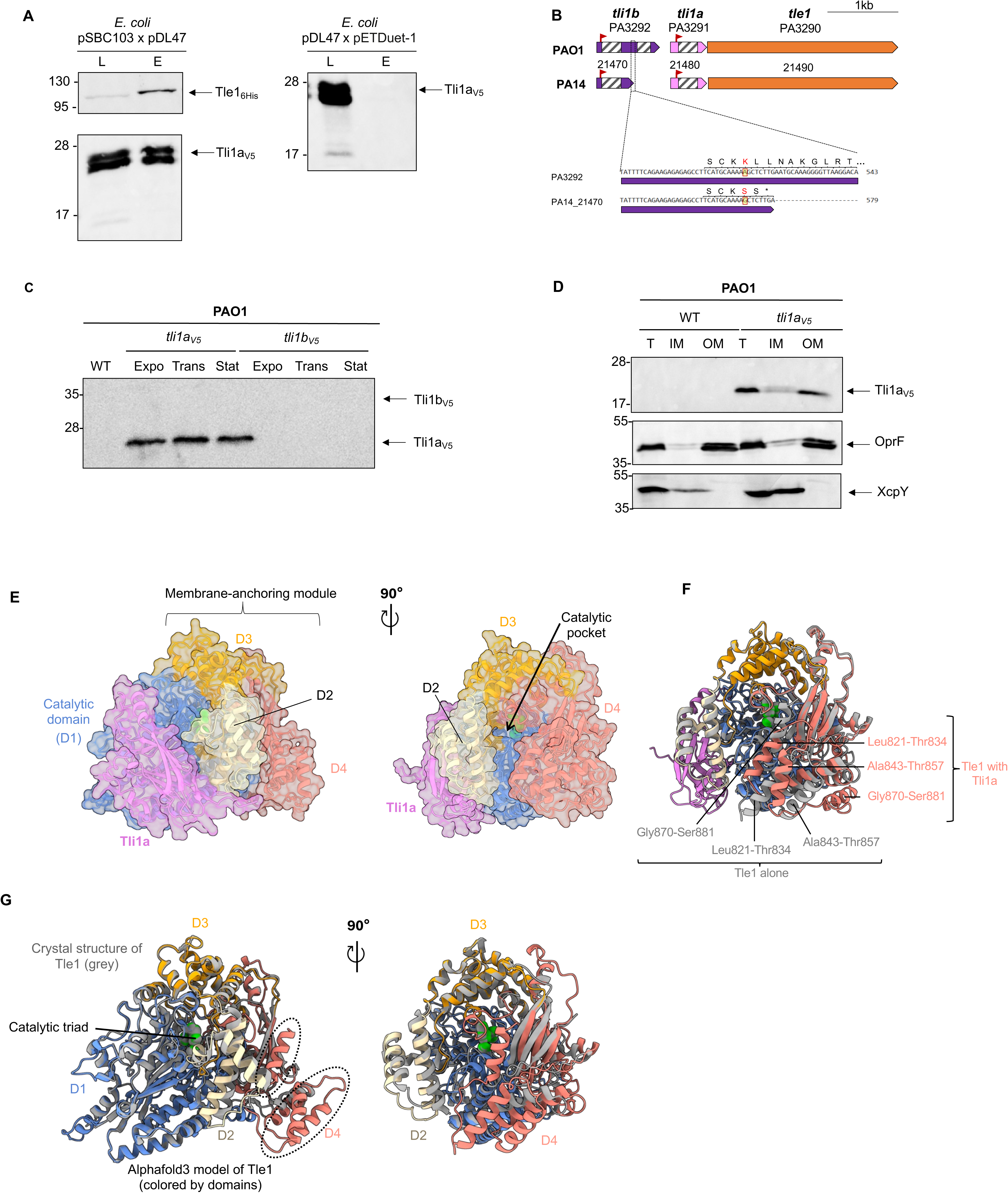
Tli1a is the immunity protein of Tle1. **(A) Tle1 interacts with Tli1a.** Co-purification assays in batch with Ni-NTA agarose resin were done using BL21 (DE3) pLysS to produce both Tle1_6His_ and Tli1a_V5_ from pSBC103 and pDL47 or only Tli1a_V5_ with pET-duet as a control. The lysate (L) and eluted (E) fractions were collected and subjected to SDS-PAGE (11%) and Western-blot analysis using anti-His antibody (upper) and anti-V5 antibody (lower). The position of the proteins and the molecular mass markers (in kDa) are indicated. **(B) Comparison of *tli1* genes and Tli1 proteins between PAO1 and PA14 strains.** The lipoprotein signal peptides are represented with red flags, the DUF3304 with striped boxes (oblique). **(C) Immunodetection of Tli1a_V5_ and Tli1b_V5_ in PAO1.** Bacteria WT or with chromosomally encoded Tli1a_V5_ or Tli1b_V5_ translational fusions were subjected to immunoblotting using the anti-V5 antibodies. The position of the proteins and the molecular mass markers (in kDa) are indicated. **(D) Localization of Tli1a_V5_ in PAO1 outer membrane.** Total membranes from PAO1 WT or with chromosomally encoded Tli1a_V5_ translational fusion were subjected to fractionation and treated with SLS to solubilize the inner membrane (IM), allowing separation from the outer membrane (OM). Anti-V5 antibodies were used for the detection of Tli1a. OprF and XcpY were used as outer membrane and inner membrane controls respectively. **(E-G) Structural model of Tle1 inhibition by Tli1a.** (E) AlphaFold 3 predicted structure of the complex Tle1 (monomer)-Tli1a (monomer) represented as transparent surface. Domains of Tle1 are colored as follows: D1, blue; D2, wheat; D3, orange; D4, red, and Tli1a is shown is pink. (F) Superimposition of the complex in (E) and the AlphaFold model of Tle1 alone (in grey). The three helices that change orientation in the complex compared to Tle1 alone in the C-terminal are indicated by a black line with their residue numbers. (G) Superimposition of the Tle1 crystal structure (in grey) and the AlphaFold 3 model of Tle1 (colored by domains).

The study of Tle1 immunity proteins continued with bioinformatics analyses. Comparison of *tle1* loci from *P. aeruginosa* strains PAO1 and PA14 revealed a difference in the annotation of the *tli1b* gene (PA3292), even though the loci were 98.6% identical (Fig. 3B top). In PA14, the gene PA14_21470 encodes a protein 95% identical to the N-terminal part of Tli1b (first 171 amino acids). Like the PA14_21480 gene homologous to *tli1a*, this gene is a candidate as Tle1^PA14^ immunity proteins because they are predicted to encode lipoproteins and contain a putative DUF3304. An alignment between the sequences of *tli1b* (PA3292) and the PA14 gene PA14_21470 suggested that the insertion of an A nucleotide at an A-rich sequence in PA3292 is responsible for a frameshift, extending protein synthesis and adding a second shorter DUF3304 (Fig. 3B bottom). A search via InterPro for proteins with a single DUF3304 domain such as Tli1a^PA01^, PA14_21470 (Tli1a^PA14^) or PA14_21480 (Tli1b^PA14^) yielded 2,360 proteins with this architecture, whereas there were only 9 proteins with two DUF3304 domains such as Tli1b^PAO1^. Given the difficulty of detecting Tli1b in *E. coli*, we speculate that the PA3292 gene is a pseudogene which has lost the ability to encode a functional protein.

To go further into Tle1 immunity proteins characterization, we chose to study them in *P. aeruginosa* and notably to determine their cellular localization. Chromosomally encoded Tli1a_V5_ and Tli1b_V5_ translational fusions were engineered in *P. aeruginosa* chromosome. While Tli1a_V5_ was readily observed in early exponential, late exponential and in stationary phases, Tli1b_V5_ was not detected under any of the growth conditions tested, supporting our previous hypothesis of a pseudogene (Fig. 3C). The cellular localization of Tli1a in *P. aeruginosa* was then determined by detergent solubilization (Fig. 3D). As expected from the presence of a lipoprotein signal sequence and the +2 and +3+4 rules, Tli1a was found in the outer membrane fraction with the control porin OprF.

The molecular mechanism underlying Tle1 inhibition by Tli1a was then investigated by predicting the structure of their complex using AlphaFold 3. We obtained a high-confidence structural model of Tle1-Tli1a complex that is supported by low PAE scores across both proteins and high ipTM (0.77) and Alphabridge (0.87) scores (Fig. 3E; Fig. S4). Consistently, Foldseek confirmed the high similarity between the experimental crystal structure of Tle1 (PDB ID: 6H3N) and the AlphaFold 3 structure, with a rsmd of 0.705 Å between 578 aligned Cα atoms (29). As previously described (17), Tle1 consists of two domains: a N-terminal catalytic phospholipase domain (residues 1-450) with an α/β-hydrolase fold, and a C-terminal membrane-anchoring domain in an open conformation (residues 451-882), subdivided into three elements (D2, D3, D4) (17). In the Tle1-Tli1a complex structure, Tli1a predominantly interacts with the catalytic domain of Tle1, and importantly without directly obstructing the catalytic pocket (Fig. 3E, compare the two orientations of the model). Notably, Tli1a engages Tle1 along its entire length, forming a continuous interface of 1912.5 Å², stabilized by hydrogen bonds and salt bridges according PDBePISA (Fig. 3E). The DUF3304 domain of Tli1a (residues 45-155) adopts a β-sandwich fold consisting of two opposing β-sheets that insert into a groove formed mainly formed by the catalytic domain of Tle1, with additional contacts to D2, confirming the BACTH data (Fig. S3). Most interactions identified by Alphabridge involve a prominent loop of Tli1a (residues 69-92) extending from its β-sandwich core, which engages residues 14-16 and 438-446 of Tle1 in D1 (piCSi : 0.85) (Fig. S4B). To assess whether Tli1a binding impacts Tle1 conformation, we superimposed the models of Tle1 alone and in complex with Tli1a on the one hand (Fig. 3F) and of the crystal structure and the model of Tle1 on the other hand (Fig. 3G). D3 is thought to function as a Tle1 cap region located above the catalytic site, the two amphipathic helices of D2 and the amphipathic β-sandwich of D4 are predicted to be inserted into the membrane (17). Interestingly AlphaFold 3 predicts structural features that are absent from the crystal structure, notably the C-terminal 69 residues of the D4 domain, which include three α-helices (Leu821-Thr834, Ala843-Thr857, Gly870-Ser881), with good confidence for the first two helices (Fig. 3G). It is noteworthy that these three helices move in the Tle1 model with Tli1a (Fig. 3F, the helices (represented in red in the model with Tli1a) have not the same orientations than in Tle1 alone (represented in grey)), suggesting that interaction with the immunity protein induces structural changes in Tle1.

The presence of Tli1a in the outer membrane of *P. aeruginosa* corroborates *in silico* prediction of an outer membrane lipoprotein and is consistent with the neutralization of Tle1, which operates from the periplasm.

### The complex of the Tle1 effector and the Tla1 chaperone with VgrG4a

To gain molecular insight into the role of Tla1 and VgrG4a during Tle1 secretion, we first confirmed the interactions between Tle1, Tla1 and VgrG4a using co-purification assays. A Strep-tagged version of the C-terminal domain of VgrG4a, and an His-tagged version of Tla1 were engineered. The recombinant proteins were coproduced in *E. coli* BL21(DE3) pLysS. The bacterial lysate was loaded to a StrepTactin matrix, and VgrG4a_CterStrep_ was eluted with desthiobiotin. Tla1_6His_ was copurified with VgrG4a_CterStrep_ only when both proteins were co-produced together (Fig. 4A, compare left and right panels). As expected for an adaptor protein, Tla1 directly interacts with Tle1. Next, interactions between VgrG4a and Tle1 were confirmed by affinity chromatography on StrepTactin matrix with VgrG4a_CterStrep_ and Tle1_6His_ (Fig. 4B) and between Tle1 and Tla1 by affinity chromatography on cobalt matrix with Tle1_6His_ and a V5-tagged version of Tla1 (Fig. 4C). Together, these experiments validate the interactions between Tla1 and VgrG4a, VgrG4a and Tle1 and Tle1 and Tla1.

**Figure 4:**
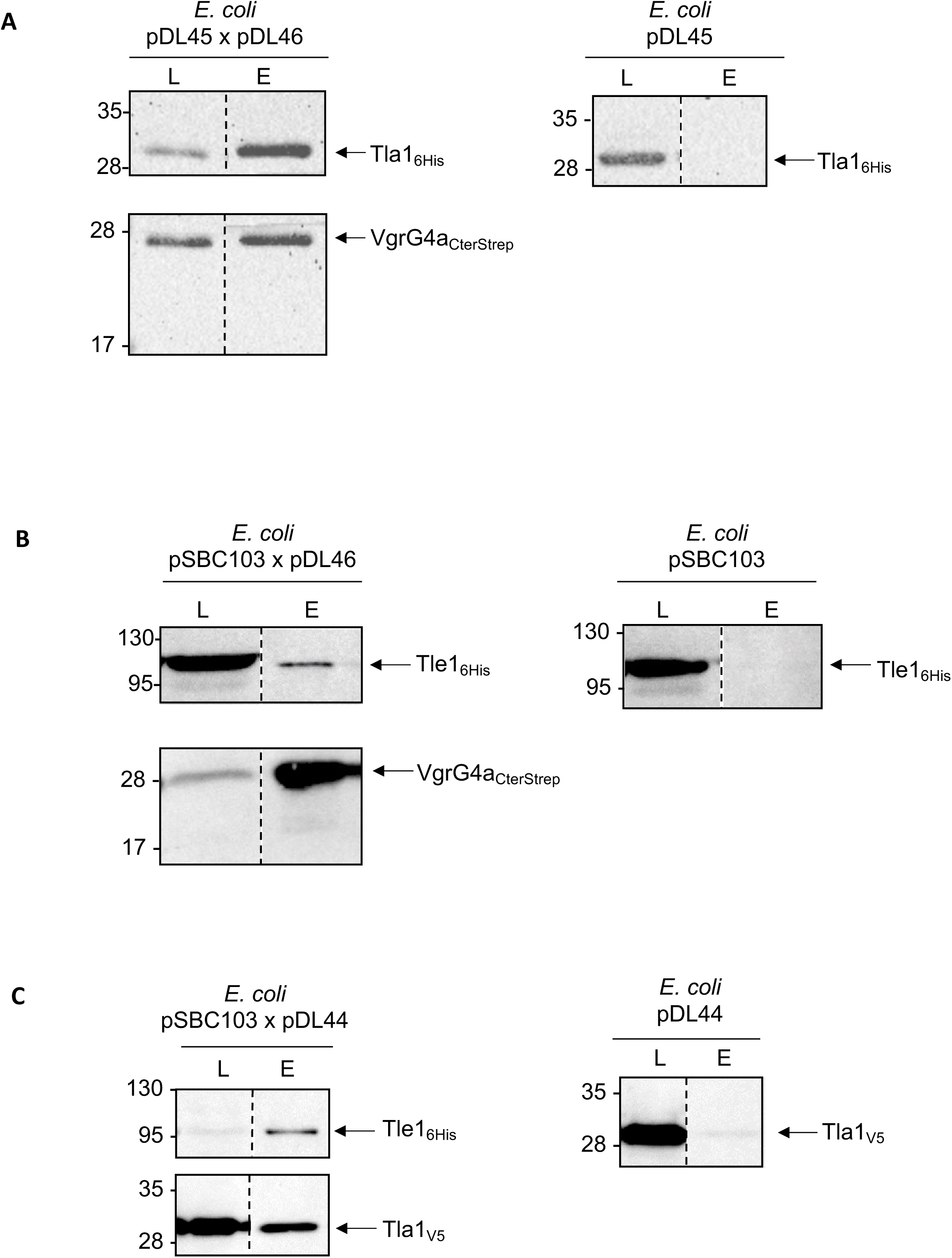
**(A) Tla1 interacts with the C-terminal domain of VgrG4a.** Co-purification assays in batch with StrepTactin resin were done using BL21(DE3) pLysS to produce both Tla1_6His_ and VgrG4a_CterStrep_ from pDL45 and pDL46 (left) or Tla1_6His_ (right). **(B) Tle1 interacts with the C-terminal domain of VgrG4a.** Co-purification assays in batch with StrepTactin resin were done using BL21(DE3) pLysS to produce both Tle1_6His_ and VgrG4a_CterStrep_ from pSBC103 and pDL46 (left) or Tle1_6His_ (right). **(C) Tle1 interacts with Tla1.** Co-purification assays in batch with cobalt resin were done using BL21(DE3) pLysS to produce both Tle1_6His_ and Tla1_V5_ from pSBC103 and pDL44 (left) or Tla1_V5_ (right). In (A), (B) and (C), the lysate (L) and eluted (E) fractions were collected and subjected to SDS-PAGE and Western-blot analyses using anti-His antibody (Upper) and anti-streptavidin or V5 antibody (Lower). The position of the proteins and the molecular mass markers (in kDa) are indicated.

Following the experimental characterization of this Tle1/VgrG4a/Tla1 interaction network, we used the AlphaFold 3 protein structure prediction program to gain insight into its structural and molecular determinants. We obtained a high-confidence model of a complex composed of a VgrG4a trimer (each protomer being annotated VgrG4a (1), VgrG4a (2) and VgrG4a (3) in figures 5A & 5B, and Fig. S5), a Tle1 monomer and a Tla1 monomer, as supported by the PAE plots, Alphabridge score (0.89) piCSi scores and pLDDT score (Fig. 5 and Fig. S5A, B, D). The trimeric VgrG4a model is similar to experimental structures of the spike of the T6SS puncturing device, such as the *P. aeruginosa* VgrG1 crystal structure (PDB ID: 6H3N) identified as a structural homolog using Foldseek (29) (rmsd of 0.94 Å between 1275 aligned Cα atoms). It contains a C-terminal extension (residues 627-688) that ends with a short α-helix (residues 672-680). The distal end of this VgrG extension (residues 668-688) interacts with Tle1 and Tla1, occupying a shallow crevice at the Tle1-Tla1 interface (piCSi: 0.76 and 0.8 respectively). Notably, VgrG4a residues 685-687 form a 3-residue β-strand that assembles into a three-stranded antiparallel β-sheet with Tla1 (indicated with a black arrow in the left insert of Fig. 5A). Noteworthy, the increase of pLDDT values in this region when bound to Tle1 and Tla1 supports the reliability of this interaction (Fig. 5B, compare the pLDDT values of the last 20 residues for each VgrG4a protomer). However, the low pLDDT score for the unstructured region of the VgrG4a C-terminal extension, located near the VgrG4a gp5-like β-prism, indicate that this flexible segment may adopt different positions with respect to the VgrG4a spike (Fig. S5A & S5D). Tla1 displays a two-domains architecture typical of DUF4123 proteins (30), with an N-terminal DUF4123 domain (residues 27-148) adopting a mixed α/β fold and a mainly α-helical C-terminal domain (Fig. S5E). The AlphaFold 3 model shows that only the catalytic domain of Tle1 interacts with Tla1 and VgrG4a.

**Figure 5:**
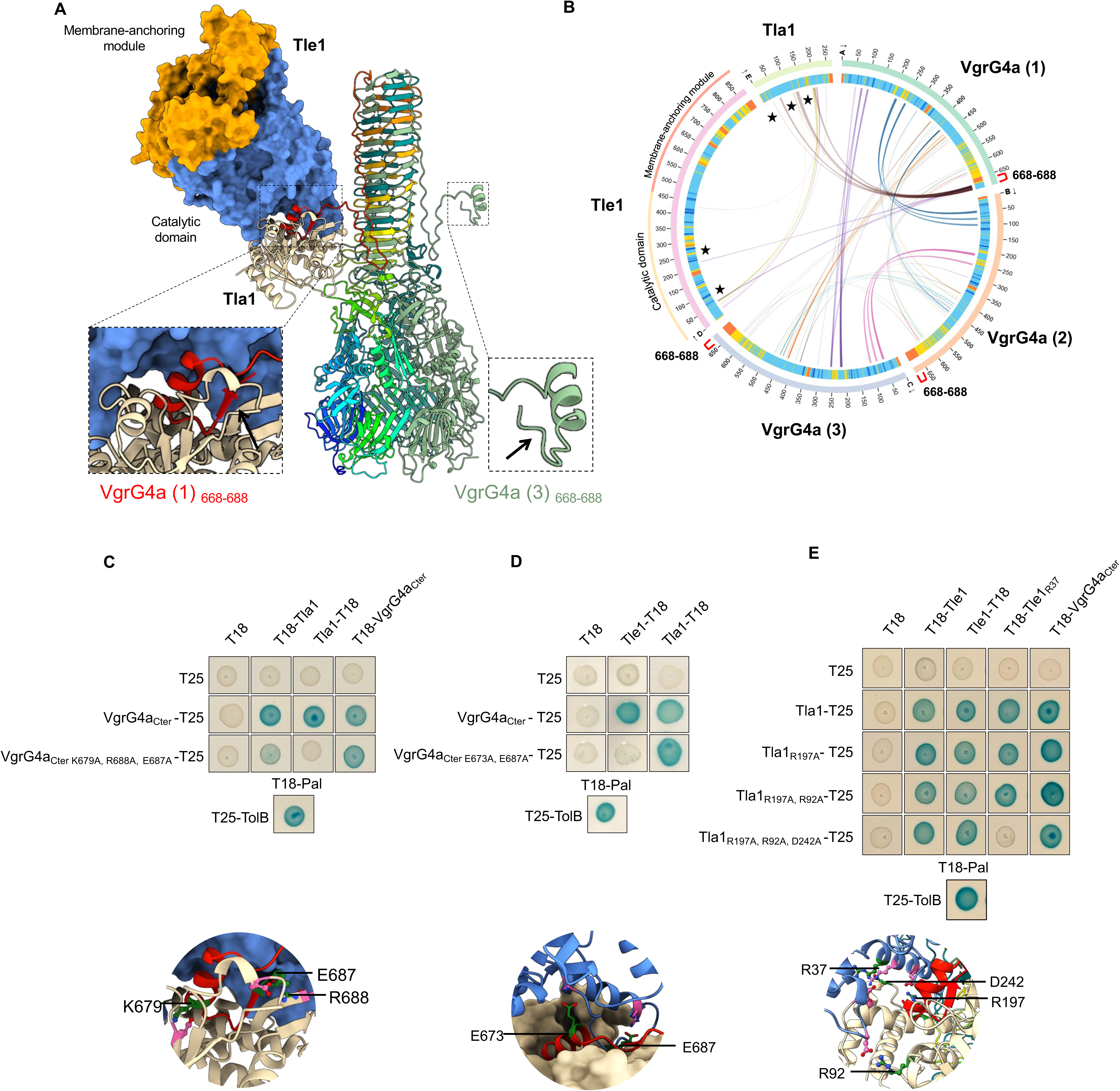
C-terminal extension of VgrG4a recruits Tle1 and its chaperone Tla1. **(A) AlphaFold 3 predicted structure of the complex between a trimer of VgrG4a, a monomer of Tle1 and a monomer of Tla1.** Inserts highlight the structured distal end of VgrG4a (protomer 1) in interaction with Tle1 and Tla1 that harbors the three-residues β-strand (685–687) (left) and the unstructured distal end of VgrG4a (protomer 3) (right). The black arrows indicate the three-residue β-strand (685–687), which is not formed in VgrG4a (3) (also absent in VgrG4a(2) but not visible here). **(B) AlphaBridge diagram of the predicted complex.** The outer and inner rings show the number of residues and the pLDDT (predicted local distance difference test) for each chain, respectively. Colors in the outer rings correspond to pLDDT confidence scores: blue (very high), cyan (high), yellow (low), and orange (very low). Regions that make contact at the VgrG4a1-Tla1 and VgrG4a1-Tle1 interfaces are indicated by curves marked with black stars. The last 20 residues of each VgrG are indicated by red square bracket on the diagram. **(C-E) Bacterial two-hybrid assays validating predicted interfaces** by alanine substitution by directed mutagenesis in *vgrG4a, tla1* and *tle1* genes cloned in BACTH vectors, targeting residues involved in salt bridges or hydrogen bonds. Substituted residues are indicated in green in the bottom inserts. **(C)** VgrG4a-Tla1 **(D)** VgrG4a-Tle1 **(E)** Tle1-Tla1. The blue color of the colony reflects interactions between two proteins. TolB and Pal are two proteins known to interact but unrelated to the T6SS. The experiment was performed in triplicate, and a representative result is shown.

We used AlphaBridge and PDBePISA to evaluate the reliability and nature of interactions assembling this VgrG4-Tle1-Tla1 complex (Fig. 5B). The last 20 residues of VgrG4a interact with the catalytic domain of Tle1 (Fig. 5B), as observed in BACTH (Fig. 2E). PDBePISA identified 16 hydrogen bonds and 10 salt bridges, including multiple salt bridges formed by VgrG4a residues E673 and E687 with Tle1 residues K61 and R47. Contacts between Tla1 and VgrG4a are primarily mediated by the N-terminal region of Tla1 containing the DUF4123 domain (residues 32-33, 60-67, 117, 136-139) and the C-terminal extension of VgrG4a (residues 669–688) (Fig. 5B). In particular, PDBePISA identified six salt bridges involving the VgrG4a residues K679 and R688 and the Tla1 D30 and E61 residues. Moreover, contacts between Tle1 and Tla1 mostly engage the Tle1 catalytic domain, as observed in BACTH (Fig. 2E), and the C-terminal part of Tla1 (residues 190-200, 207-210, 243) (piCSi: 0,82-0,86). Additionally, four residues of the Tla1 DUF4123 domain (R92, R93, D35, N120) make contacts with Tle1 (D429, R107 and V53). This large 1070 Å^2^ Tle1-Tla1 interface is stabilized by hydrogen bonds and salt bridges.

To experimentally validate the relevance of these protein-protein contacts assembling the VgrG-Tle1-Tla1 complex, we performed alanine substitution by directed mutagenesis approach on *tle1*, *tla1* and *vgrG4a* genes cloned in BACTH vectors, targeting mostly residues involved in salt bridges. As shown in figure 5C, the interaction between Tla1 and VgrG4a was lost when three residues (K679 and R688 forming salt bridges, and E687 involved in hydrogen bonding), were successively substituted for an alanine in VgrG4a_Cter_. This VgrG4a_Cter K679A, R688A, E687A_ variant was functional since it is still able to interact with VgrG4a_Cter_ (Fig. 5C). The interaction between VgrG4a and Tle1 was lost when two residues were changed in VgrG4a_Cter_, E673 and E687 (Fig. 5D). This VgrG4a_Cter E673A, E687A_ was still able to bind Tla1. Lastly, we confirmed that four residues, one in Tle1, R37, and three in Tla1, R197, R92, D242, all involved in salt bridges, are involved in the interaction between the toxin and its chaperone (Fig. 5E). Importantly, these Tle1 and Tla1 variants retained binding to VgrG4a_Cter_, confirming that the mutations specifically disrupted the Tle1-Tla1 interface rather than overall protein folding. In conclusion, the three predicted interaction interfaces, Tle1-Tla1, Tla1-VgrG4a, and Tle1-VgrG4a, were validated *in vivo*.

## Discussion

Our findings establish Tle1 as a bactericidal toxin of *P. aeruginosa* and uncover the molecular bases governing its delivery and neutralization. By integrating AlphaFold-based structure prediction with complementary protein-protein interaction assays, we delineated the specific interplay between Tle1 and its immunity protein Tli1a on the one hand, and between Tle1, the adaptor/chaperone Tla1 and the structural component VgrG4a on the other hand. This work provides a mechanistic framework for understanding how Tle1 is deployed through the T6SS while being tightly controlled to avoid self-intoxication.

We demonstrated that Tle1 exerts a bactericidal effect, although moderate, when targeted to the periplasm from where it inserts into the membrane (Fig. 1). This finding is consistent with previous reports describing Tle1 family effectors as antibacterial phospholipases with variable toxicity in *Burkholderia thailandensis*, entero-aggregative *E. coli* (EAEC), *Klebsiella pneumoniae*, *Aeromonas hydrophila*, *V. cholerae*, *B. cenocepacia* and *Xanthomonas oryzae* (10, 17, 23–25, 31–36). For instance, Tle1^BT^ has been shown to induce membrane permeability, as demonstrated by the entry of propidium iodide into *B. thailandensis* cells lacking the Tli1^BT^ immunity protein (10). Similarly, Tle1^VC^ and Tle1^KP^ increase the permeability of target bacteria (24, 37). Following the action of Tle1^KP^, a membrane depolarization effect has been observed (31). In line with our observation, intra-species competition macrocolony assays in *P. aeruginosa* showed that Tle1 is not a highly active toxin, even in derepressed backgrounds for T6SS expression used in that study (38). While Tle3, a previously characterized Tle of *P. aeruginosa* (15), was active in both *retS* and *rsmA* mutants’ backgrounds, a slight Tle1 antibacterial activity was only observed in *retS*, the double *rsmA tle1* mutant exhibiting a *rsmA* profile. The individual impact of toxins is thus highly variable in *P. aeruginosa*. The moderate effect observed here may reflect the intrinsic catalytic properties of Tle1^PA^.

Two genes encoding putative Tle1 immunity proteins, *tli1a* and *tli1b*, are present upstream of *tle1*. In agreement with its role as an immunity protein, Tli1a was shown to interact directly with Tle1 by BACTH and pull-down assays (Fig. 2 and 3). Furthermore, Tli1a has been observed in the outer membrane of *P. aeruginosa* (Fig. 3D), a localization that would enable it to counteract the toxicity of Tle1 during a fratricidal attack. This localization is consistent with the other previously characterized immunities of Tle1 in various bacteria (25). As we had previously done for Tle3 (15), we tried to demonstrate the neutralization of Tle1 in *E. coli* by Tli1a. To achieve this, we attempted to co-express *tli1a* and ss-*tle1* (Fig. Sup 6A). However, this was unsuccessful and SS-tle1 was still toxic (Fig. S6, compare line 3 with 4). Although we were unable to observe the production of Tli1b in *P. aeruginosa* (Fig. 3C), the other putative immunity of Tle1, we also attempted, without further success, the co-overproduction of Tle1 with Tli1b (Fig. S6B) or with both putative immunities in *E. coli*, from the same transcript under the same promoter by cloning *tli1a, tli1b* and *ss-tle1* in tandem (Fig. S6B). Tli1a was produced, whereas Tli1b was not (Fig. S6C) and SS-tle1 was still toxic. However, one piece of data from the literature is consistent with our result of an immunity role for Tli1a. Studying intra-species competition by a macrocolony assay, Rudzite *et al.* (38) were able to delete the *tli1a* and *tli1b* genes by also deleting *tle1*. Since the genes encoding immunity are essential genes, the viability of this mutant supports a role for Tli1a as an immunity protein of Tle1. Regarding Tli1b, analysis of the conservation of this gene in other strains of *P. aeruginosa* combined with its unusual architecture, consisting of two DUF3304 domains (one complete and one shorter), suggests that it may be a pseudogene. The *tli1b* gene is predicted in an operon structure with *vgrG4a* and *tla1* in PAO1 (Pseudomonas.com). If it is indeed transcribed, we were unable to detect the corresponding protein in *P. aeruginosa* (Fig. 3C), despite the presence of a RBS upstream of its ATG. This genetic organization is reminiscent of that of *tle1* in EAEC, near which there are two genes encoding potential immunity proteins. Neutralization of Tle1^EAEC^ by Tli1 is sufficient, while the second gene *tli1b* carries a mutation resulting in a truncated protein (25). The gene encoding the Tle1^VC^ protein of *V. cholerae* is also located near two genes encoding putative immunity proteins, with the same DUF2931 domain. However, while Tli^Tox−^ is sufficient for protection against Tle1^VC^, Tli^Tox+^ is not, despite sharing 80.24% sequence identity (31).

Structural predictions using AlphaFold 3 generated high-confidence models indicating that Tli1a binds the catalytic domain of Tle1 without occluding the substrate entry pocket (Fig. 3E-G). Strikingly, unlike Tli3 from adherent-invasive *Escherichia coli* (AIEC), which inserts into the catalytic crevice of Tle3 (39), none of the AlphaFold 3 models suggest that Tli1a enters the catalytic pocket, but rather they predict interactions outside of it. One hypothesis could be that Tli1a prevents or disturbs the insertion of Tle1 into the membrane where it exerts its activity. To test this, QMEANBrane was used to model the membrane insertion of Tle1 alone or in the presence of Tli1a. Interestingly the Tle1 membrane insertion model shows that the two amphipathic helices in D2 insert into the membrane (Fig. S7A), D2 outlined in red dotted line), but no longer in the presence of Tli1a (Fig. S7B). Moreover, the amphipathic β-sandwich in domain D4 shows reduced insertion into the bilayer. In agreement with this hypothesis, conformational changes in the helices of Tle1 D4 domain were observed in the model with Tli1a (Fig. 3E-F). This positioning, together with the demonstration that Tli1a is an outer membrane lipoprotein (Fig. 3D), could also suggest a mechanism in which Tli1a neutralizes Tle1 from the periplasmic side, thereby preventing its correct membrane insertion and access to its phospholipid substrate. As a complementary mechanism, the interaction between Tli1a and Tle1 at the outer membrane could also act as a spatial sequestration mechanism, preventing Tle1 from reaching and degrading the inner membrane, which is assumed to be the primary target of Tle. Two original mechanisms for neutralizing Tle in *P. aeruginosa* have been described so far. Indeed, the Tli4 immunity protein counteracts its cognate TplE (Tle4) toxin through a crab-claw neutralization mechanism (40). PldA (Tle5a) and PldB (Tle5b) also interact with their cognate immunity proteins outside the catalytic pocket by a cup-saucer mechanism. In this case, there would be large contact interfaces between the toxin and its cognate immune proteins, which sequester it structurally without necessarily directly blocking the active site. The “saucer” made by immunity proteins partially encompasses the catalytic domain of PldB (13). For Tli5a, the immunity of PldA (Tle5a), a conserved region of 25 amino acids, a β10-β12 domain, has been described (41), and for the three Tli5b immunity proteins of PldB (Tle5b), Sel1-like repeats (SLRs) have been identified (13, 42). The presence of these two motifs was not detected in Tli1a. Finally, a last example is the DNA binding site of the nuclease effector Tde1 from *Agrobacterium tumefaciens* that becomes disordered when bound to its immunity protein, demonstrating that effector neutralization is not limited to restricting access to the active site (43).

Protein-protein interaction assays revealed a network of interactions within *tle1* locus. Specifically, we detected interactions between Tle1 and its adaptor Tla1, as well as between Tla1, Tle1 and the structural component VgrG4a (Fig. 2 & 5). These data support a model in which Tle1 is loaded onto the distal end of the type VI secretion spike through Tla1, in line with adaptor-mediated effector recruitment described for other T6SS toxins (44, 45). Structural predictions using AlphaFold 3 provided high-confidence models of a T6SS puncturing device in which a VgrG4a trimer accommodates Tle1 *via* Tla1. Due to computational constraints, we could not model a complex with three Tle1, three Tla1 monomers, and full-length VgrG4a. However, when using truncated VgrG4a containing only the C-terminal extensions (Fig. S8), AlphaFold 3 predicted that each extension could recruit one Tle1-Tla1 complex, leading to the formation of the three-residue β-strand by the last VgrG4a residues (685–687), as observed in Fig. 5A. It is also conceivable that each VgrG4a protomer may interact with a distinct effector, or alternatively, one or two remain unbound. Indeed, VgrGs involved in the secretion of several toxins have been described (46, 47), but it is not formally known whether this occurs simultaneously. In addition, the identification of VgrG heterotrimers (48) calls into question the assumption that VgrG4a forms only a homotrimer. However, at this stage, no other partners (effectors, VgrG or PAAR) of VgrG4a have been identified.

A helix-turn-helix region in the C-terminal end of VgrG6 and VgrG14 of *P. aeruginosa* was modeled as being in contact with the Tap6 and Tap14 adaptors, through their N-terminus containing the DUF4123 (30). The C-terminal domain of adaptors was shown to mediate the interaction with cognate effector (30). Despite the absence of the helix-turn-helix region in VgrG4a, our model agrees with this data. Tla1 has been observed to interact with its N-terminus containing the DUF4123 with the C-terminus of VgrG and with its C-terminus with Tle1 (Fig. 5). Remarkably, the predicted interfaces between Tle1, Tla1, and VgrG4a were validated experimentally *in vivo*, underscoring the reliability of recent AI-based structural predictions for modeling secretion system complexes. Such integrative approaches are increasingly used to bridge the gap between *in silico* models and experimental validation (49).

Finally, it was observed that Tle1 can interact in the periplasm with its immunity protein Tli1a and in the cytoplasm with its adaptor Tla1 and VgrG4a, forming a ternary complex. One could discuss the differences in affinity of these different interactions: the interaction with the immunity protein must be stronger and more stable in order to inhibit the toxicity of Tle1, whereas Tla1 must detach from the toxin at the moment of its secretion, just as VgrG4a must release it so that it can fulfill its function in the target bacterium. Do adaptors really dissociate? Do VgrGs really release their effectors into target cells? So many open questions, which constitute a working model of the Tle1 secretion process, which offers exciting prospects for future research.

Altogether, our results extend the current understanding of Tle family toxins by providing a structural and functional framework for Tle1 activity in *P. aeruginosa*. They highlight the interplay between effector, adaptor, structural proteins, and immunity proteins, which collectively ensure efficient toxin delivery while preventing self-intoxication.

## Experimental procedures

### Bacterial strains, growth conditions, and plasmid construction

All strains of *P. aeruginosa* and *E. coli* used in this study are listed in Table 1. In brief, the *E. coli* strains CC118λPir and K-12 DH5α were used for cloning procedures, while BL21(DE3) pLysS was employed for protein expression/production under T7 promoter, and BTH101 for BACTH assays. Cultures were grown in LB medium or TSB medium (for *P. aeruginosa*) at 37°C or 30°C, with specific growth conditions provided in the main text when required. Plasmids were introduced into *P. aeruginosa* through triparental mating, facilitated by the helper plasmid pRK2013 (Table S1). Plasmid maintenance was ensured by supplementing media with appropriate antibiotics: ampicillin (50 μg/ml) for *E. coli*, kanamycin (50 μg/ml) for *E. coli*, chloramphenicol (30 μg/ml) for *E. coli*, streptomycin (30 μg/ml for *E. coli* and 2,000 μg/ml for *P. aeruginosa*). Cloning was performed by sequence and ligation independent cloning (SLIC) (50) and cloned sequences were confirmed by DNA sequencing (Eurofins). Table 1 provides a list of plasmids constructed and used, and the oligonucleotides synthesized by Eurogentec and IDT.

### Heterologous toxicity assays

*E. coli* BL21(DE3) pLysS strains harboring plasmids producing either cytoplasmic or periplasmic targeted proteins were cultured overnight at 37°C in LB medium supplemented with 0.4% glucose. Serially diluted bacterial suspensions (10 μl) were spotted onto LB agar plates containing either 0.1 mM IPTG or 0.4% glucose and incubated at 37°C for 16 hours as described in (15).

### Bacteriostatic or bactericidal effect of the toxin

This assay has been done as in (17). From stationary-phase overnight cultures, fresh LB medium containing 1% glucose was inoculated to an OD_600_ of 0.1. Bacteria were cultivated at 37°C to OD_600_ = 0.6 in LB medium containing 1% glucose, washed twice with LB before induction with 0.1 mM IPTG. At time 30- and 60-min post-induction, an aliquot was recovered and chilled in ice water for 2 min. Cells were pelleted at 6,000 X *g* at 4°C and re-suspended in ice-cold fresh LB. After normalization to an OD_600_ of 0.5, serial dilutions were done in sterile PBS and spotted on LB agar plates containing appropriate antibiotics and 1% glucose.

### Cloning procedures for *P. aeruginosa*

To generate chromosomally encoded Tli1a_V5_ and Tli1b_V5_ translational fusions in *P. aeruginosa*, primers were designed to introduce a V5 epitope sequence at the C-terminus of each target gene. Regions of 500 bp upstream and downstream of each gene were PCR-amplified using Q5 high-fidelity DNA polymerase (NEB) and the V5-tagged primers listed in Table 1. The PCR fragments were cloned into the pKNG101 suicide vector using one-step sequence- and ligation-independent cloning (SLIC) (50). After sequence verification, the resulting pKNG101 constructs were introduced into *E. coli* CC118λpir and subsequently transferred into *P. aeruginosa* via conjugation as previously described in (14). Mutants generated through double homologous recombination were screened and validated by PCR.

### Bacterial two-hybrid assay

The adenylate cyclase-based two-hybrid (BACTH) assay was employed to study protein–protein interactions, using established methods (51, 52) and the protocol previously described (15).

### Site-directed mutagenesis

Targeted mutagenesis was performed on plasmid constructs used in the BACTH system (Table 1) to assess the role of specific residues at protein–protein interfaces predicted by AlphaFold 3 (53). Salt bridges and hydrogen bonds identified by PDBePISA (48, https://www.ebi.ac.uk/pdbe/pisa/pistart.html)) were selected based on their localization and number of contacts. Mutagenic primers encoding the desired substitutions to alanine were designed following the QuickChange method. PCR amplification was carried out using a high-fidelity DNA polymerase, and parental plasmids were digested with DpnI to remove methylated templates. The resulting PCR products were transformed into *E. coli* DH5α. Positive clones were selected on LB agar containing the appropriate antibiotics, and plasmids were purified using standard miniprep procedures. All mutations were confirmed by DNA sequencing (Eurofins). Finally, the mutated plasmids were co-transformed into the BACTH reporter strain *E. coli* BTH101 and subjected to the BACTH assay.

### SDS-PAGE and Western-Blot

Protein samples corresponding to equivalent culture densities (measured by optical density at 600 nm) were resuspended in loading buffer, boiled, and subjected to SDS-PAGE. Proteins were subsequently detected via immunoblotting, following the method described by (55), using primary monoclonal antibodies against His6 (Penta His, Qiagen, 1:1,000), V5 (Bethyl Laboratories, 1:5,000), Strep (IBA StrepMAB Classic, 1:2,500), EF-Tu (Hycult-biotech, 1:20,000), XcpY (laboratory collection, 1:1,000), and Pal (laboratory collection, 1:5000). Peroxidase-conjugated anti-Rabbit IgGs or anti-Mouse (Sigma, dilution 1:5000) were employed as secondary antibodies. Protein revelation was carried out using a homemade enhanced chemiluminescence and membrane were analyzed with ImageQuant LAS4000 software (GE Healthcare Life Sciences). Protein samples corresponding to 0.1 OD_600_ units were loaded for whole-cell and spheroplast analyses, while 0.2 OD_600_ units were used for cytoplasmic, periplasmic and membranes fractions.

### Protein purification by affinity chromatography

*E. coli* BL21(DE3) pLysS cells harboring pRSFDuet-1 or pETDuet-1 derivative plasmids were cultivated in LB medium at 37°C until an OD_600_ of 0.6 was reached. The expression of the PA3290, PA3291, PA3293, or PA3294 genes was induced by adding 1 mM IPTG, followed by incubation for 2 hours at 37°C. Cells were then collected by centrifugation at 1,914 X *g* for 15 minutes at 4°C. The resulting cell pellets were resuspended in a buffer containing 50 mM Tris-HCl (pH 8.0), 300 mM NaCl, 0.5 mg/ml lysozyme, and 1 mM EDTA, and stored at −80°C.

Before lysis, the cell suspension was supplemented with 20 μg/ml DNase (SIGMA), 20 mM MgCl_2_, 1 mg/ml de lysozyme and 1 mM phenylmethylsulfonyl fluoride (PMSF). Cells were lysed by sonication, and the lysates were clarified by centrifugation at 16,000 X *g* for 30 minutes. The clarified supernatant was incubated for 3 hours with either Ni-NTA resin (Macherey-Nagel) pre-equilibrated in binding buffer (50 mM Tris–HCl pH 8.0, 300 mM NaCl, 10 mM imidazole), Strep-Tactin resin (IBA Lifesciences) pre-equilibrated in its respective binding buffer (100 mM Tris–HCl pH 8.0, 150 mM NaCl), or a Cobalt resin (Thermo Scientific) pre-equilibrated in binding buffer (50 mM Tris–HCl pH 8.0, 150 mM NaCl, 10 mM imidazole, 10% glycerol, 0,1% Triton X-100 (Sigma)).

The resins were washed with their respective washing buffers: for Ni-NTA, 50 mM Tris–HCl (pH 8.0), 300 mM NaCl, and 20 mM imidazole; for Strep-Tactin and cobalt resins, binding buffer. Target proteins were eluted using the corresponding elution buffers: binding buffer containing 500 mM imidazole for Ni-NTA for cobalt resin, binding buffer containing 5mM desthiobiotin for Strep-Tactin.

### Fractionation

Fractionation of *E. coli* cells into spheroplasts (cytoplasm and membranes) and periplasmic fractions were done as described previously (56). Outer membrane and inner membrane fractionation of *P. aeruginosa* was performed as described in (57), with a treatment of the total membrane pellet with 0.5% sodium *N*-lauroyl sarcosinate (SLS; Sigma Aldrich). Resulting membrane pellets were resuspended in SDS-PAGE sample buffer for analysis.

### Protein–protein interaction *in silico* analyses

To predict the 3D structures of protein complexes, the amino acid sequences of the proteins of interest, obtained in FASTA format from *Pseudomonas.com*, were submitted to AlphaFold 3 (https://alphafoldserver.com/welcome) (53). These predictions were produced with predicted aligned errors (PAE) and pLDDT values as confidence scores. The AlphaFold-predicted structures were subsequently analyzed with AlphaBridge (58), investigate residue-residue interactions at the predicted interfaces. Confidence in these predicted interfaces was assessed using the predicted interaction confidence score (piCS), with a default cut-off of 0,7. Finally, we used PDBePISA to analyze protein-protein interfaces of the predicted complexes (54) and the Foldseek server to identify the closest structural homologues in the Protein Data Bank. Visual representations of the structures were prepared with ChimeraX (59). QMEANBrane (https://swissmodel.expasy.org/qmean/) was used to predict membrane protein models in their naturally occurring state.

## Data availability

The data described in this study are contained within the manuscript. Coordinates of predicted structures will be deposited in the open data repository Zenodo.

## Supporting information

This article contains supporting information.

## Acknowledgments

We thank members of Bleves-Ize team and members of the LISM, in particular C. Sebban-Kreuzer and S. Canaan, for helpful discussions and support. We are grateful to M. Ba, I. Bringer, A. Brun and M. Guilbert for technical assistance. We acknowledge UCSF ChimeraX for molecular graphics that is developed by the Resource for Biocomputing, Visualization, and Informatics at the University of California, San Francisco, with support from National Institutes of Health R01-GM129325 and the Office of Cyber Infrastructure and Computational Biology, National Institute of Allergy and Infectious Diseases. The English language was improved with the assistance of AI (DeepL and ChatGPT).

## Author contributions

DL, AG, BI and SB designed and conceived the experiments. DL, CS, LS and BI performed the experiments. SB supervised the execution of the experiments. DL, CS, LS, AG, BI, and SB analyzed and discussed the data. SB wrote the paper with contribution from DL and review/editing from all the authors.

## Funding and additional information

This work was supported by the Aix-Marseille Université (amU), the Centre National de la Recherche Scientifique (CNRS), a grant from the Agence Nationale de la Recherche (ANR, ANR-21-CE11-0028) and a grant from Vaincre La Mucoviscidose (VLM, RF20240503476/1/1/69). DL was supported with a PhD fellowship from the French Research Ministry, and by the Fondation pour la Recherche Médicale, « grant number FDT202404018453 » for a 4^th^ year.

