## supplemental figures for "Molecular Insights into the bactericidal Toxin Tle1 of *Pseudomonas aeruginosa*: Interaction with VgrG, its adaptor, and immunity protein"

**Supporting information Figure 1**

**A**

**Sequence**
**Prediction:** Lipoprotein signal peptide (Sec/SPII)

Cleavage site between pos. 30 and 31. Probability 0.829392

| Protein type | Other | Signal Peptide (Sec/SPI) | Lipoprotein signal peptide (Sec/SPII) | TAT signal peptide (Tat/SPI) | TAT Lipoprotein signal peptide (Tat/SPII) | Pilin-like signal peptide (Sec/SPIII) |
|---|---|---|---|---|---|---|
| **Likelihood** | 0.0002 | 0.1727 | 0.8271 | 0 | 0 | 0 |

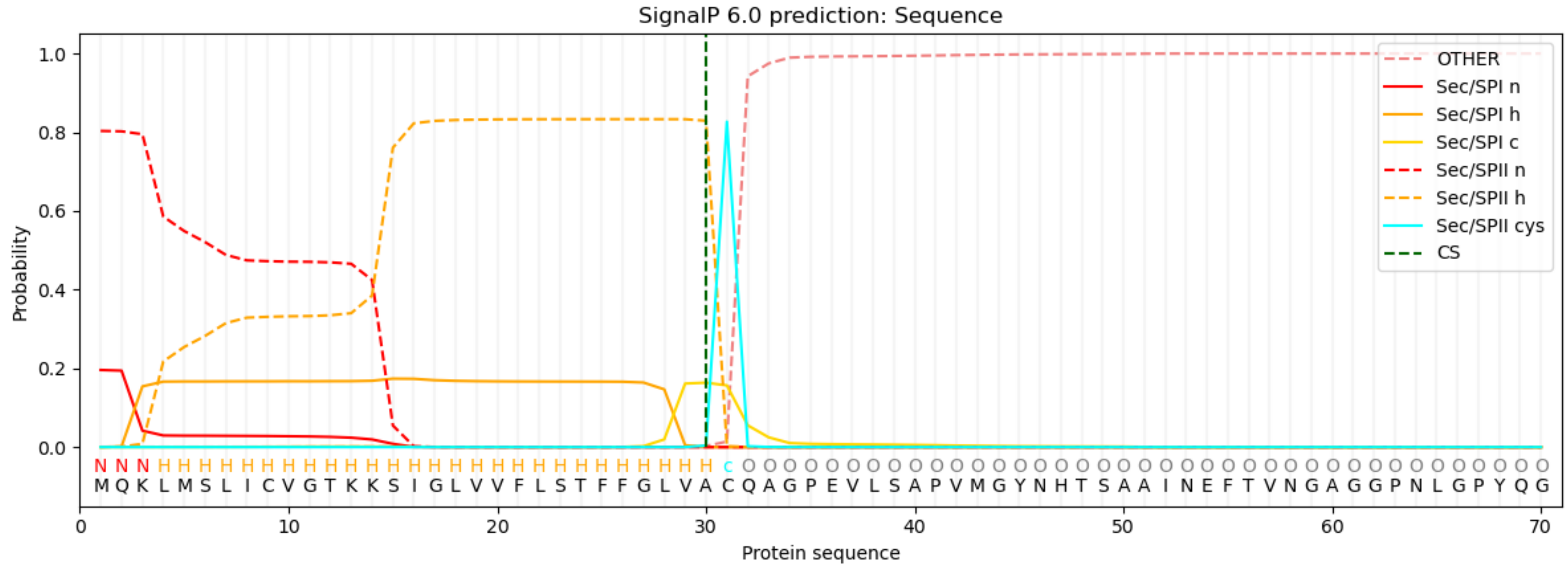


**B**

**Sequence**
**Prediction:** Lipoprotein signal peptide (Sec/SPII)

Cleavage site between pos. 20 and 21. Probability 0.994805

| Protein type | Other | Signal Peptide (Sec/SPI) | Lipoprotein signal peptide (Sec/SPII) | TAT signal peptide (Tat/SPI) | TAT Lipoprotein signal peptide (Tat/SPII) | Pilin-like signal peptide (Sec/SPIII) |
|---|---|---|---|---|---|---|
| **Likelihood** | 0 | 0 | 1 | 0 | 0 | 0 |

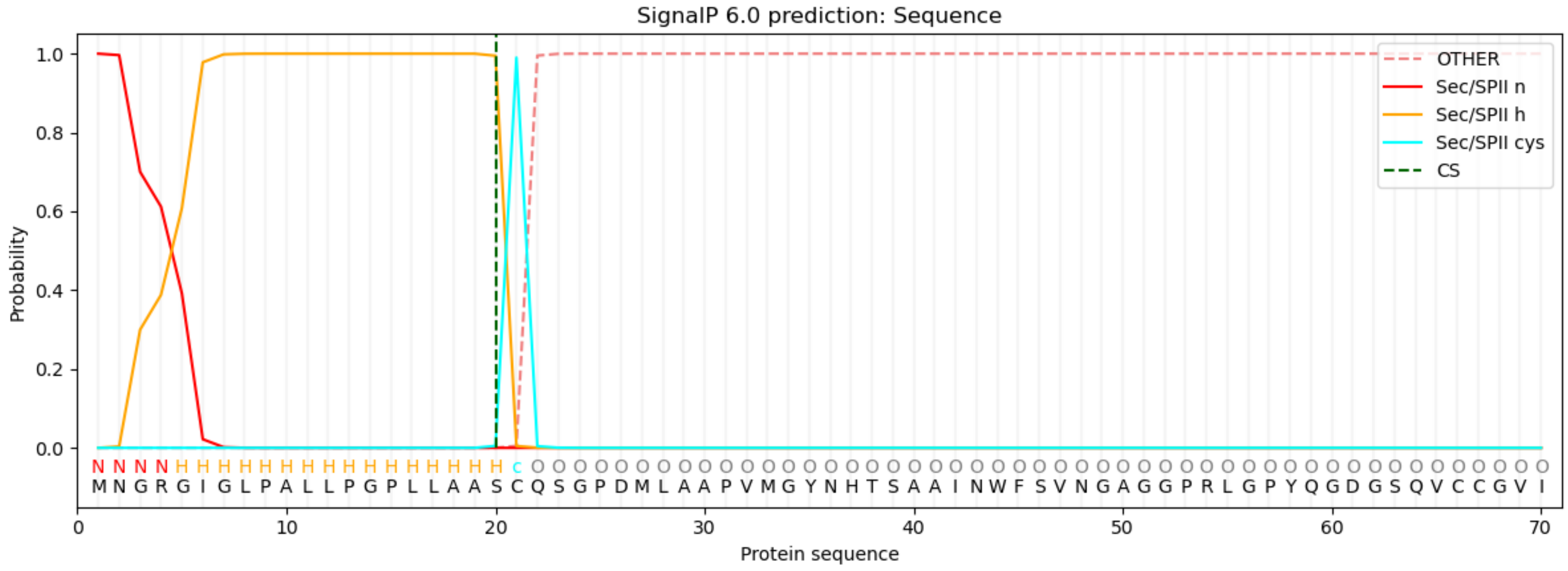


**Fig. S1: SignalP 6.0 predictions for (A) PA3291 (Tli1a) and (B) PA3292 (Tli1b).** SignalP 6.0 was used to predict a signal peptide at the N-terminus of the two corresponding proteins and the class (Sec, Tat or Lipoprotein) of this signal peptide.

### Supporting information Figure 2

**A**

```
Query: PA3291-Tli1a ID: lcl|Query_1658233(amino acid) Length: 184
Subject:PA3292-Tli1b ID: lcl|Query_1658235(amino acid) Length: 285

Clusters producing significant alignments:
<@acc_hd@><@sciname_hd@><@comname_hd@><@taxid_hd@>

Cluster:           Query_1658235     PA3292-Tli1b
Num Members:       0
Num Taxa:          0
Scientific Name:
Common Name :
Taxid:             0
Highest Bit Score: 256
Total Bit Score:   367
Percent Coverage:  92%   %
Evalue:            3e-91
Percent Identity:  75.33  %
Accession Length:  285

0            cluster member(s):
<@clust_mem_rows@>

Alignments:

>PA3292-Tli1b
Sequence ID: Query_1658235 Length: 285
Range 1: 20 to 169
Score:256 bits(653), Expect:3e-91,
Method:Compositional matrix adjust.,
Identities:113/150(75%), Positives:132/150(88%), Gaps:0/150(0%)

Query  30   ACQAGPEVLSAPVMGYNHTSAAINEFTVNGAGGPNLGPYQGDGSQVCCGVIPKRWNPNLK  89
            +CQ+GP++L+APVMGYNHTSAAIN F+VNGAGGP LGPYQGDGSQVCCGVIPK+WNPNLK
Sbjct  20   SCQSGPDMLAAPVMGYNHTSAAINWFSVNGAGGPRLGPYQGDGSQVCCGVIPKKWNPNLK  79

Query  90   VIVEWEKDPNPRAVIKRDKYGRLDESDYLRHASSYTRHKATVNVPRYDEKVCLLQVHFLP  149
             +VEWEKDP P A I+RDKYGRLD+ DYLRHASSYTRHK  V++PRY EK+CLLQVHFLP
Sbjct  80   AVVEWEKDPKPHAAIRRDKYGRLDKDDYLRHASSYTRHKMIVDIPRYSEKICLLQVHFLP  139

Query  150  CDEVAVSTTCYGPNHPKYPDKAYFEMRKST  179
            CD+VAVSTT Y   HP+YP + YF+ R+ +
Sbjct  140  CDQVAVSTTYYSQGHPEYPVRKYFQKREPS  169


Range 2: 178 to 269

Score:111 bits(277), Expect:1e-34,
Method:Compositional matrix adjust.,
Identities:52/92(57%), Positives:62/92(67%), Gaps:0/92(0%)

Query  11   GTKKSIGLVVFLSTFFGLVACQAGPEVLSAPVMGYNHTSAAINEFTVNGAGGPNLGPYQG  70
            G +    L+  L  F  L ACQAG +++S P+ GYNHTSAAIN F+VNGAGG NLGP+QG
Sbjct  178  GLRTPDWLIALLIGFLFLQACQAGSDMVSTPIGGYNHTSAAINRFSVNGAGGLNLGPFQG  237

Query  71   DGSQVCCGVIPKRWNPNLKVIVEWEKDPNPRA  102
               QVCCGV+P+ W   LK  VEWE  P P A
Sbjct  238  GAGQVCCGVVPRVWKSGLKATVEWEVVPEPGA  269
```

**B**

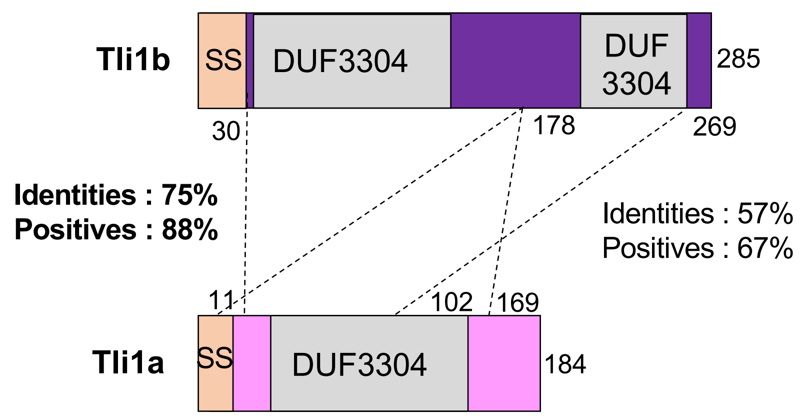


**Fig. S2 : BLAST alignment between PA3291 (Tli1a) and PA3292 (Tli1b),** showing the BLAST output **(A).** The red rectangles highlight the percentage of identity, either for the full-length proteins or for specific regions. **(B)** Schematic representation of the BLAST alignments of Tli1a (pink) and Tli1b (purple). The lipoprotein signal sequences (SS) of Tli1a (PA3291) and Tli1b (PA3292) are indicated by orange boxes. The DUF3304 domains are shown in grey and delineated; Tli1b contains two DUF3304 domains (residues 35–145 and 212–275), while Tli1a contains a single DUF3304 domain (residues 45–155). Similar regions between Tli1a and Tli1b, identified by BLAST, are indicated by dotted lines, along with their percentage of identity and the positively conserved residues with their respective residue numbers.

**Supporting information Figure 3**

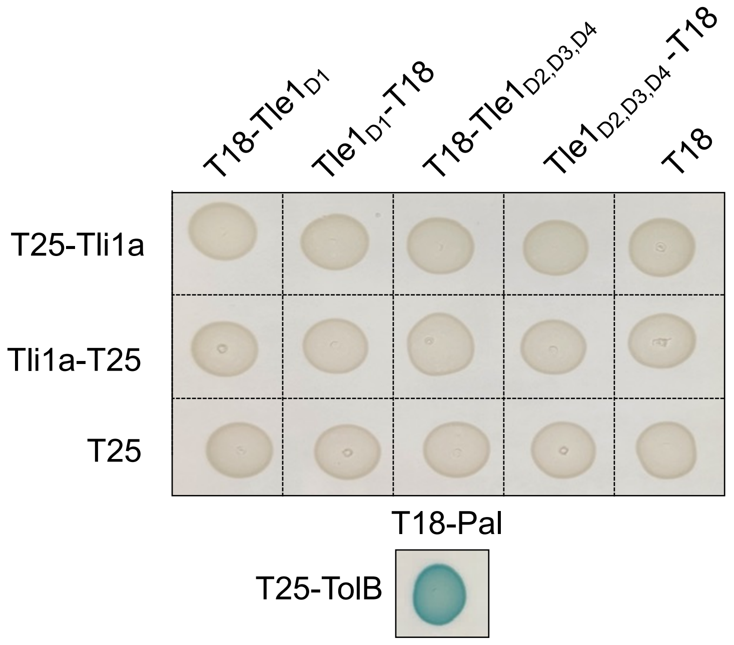


**Fig. S3 : Bacterial two-hybrid assay.** BTH101 reporter cells producing the indicated proteins or domains fused to the T18 or T25 domain of the *Bordetella pertussis* adenylate cyclase were spotted on X-gal indicator plates. The blue color of the colony reflects the interaction between the two proteins. TolB and Pal are two proteins known to interact but unrelated to the T6SS. The experiment was performed in triplicate and a representative result is shown.

**Supporting information Figure 4**

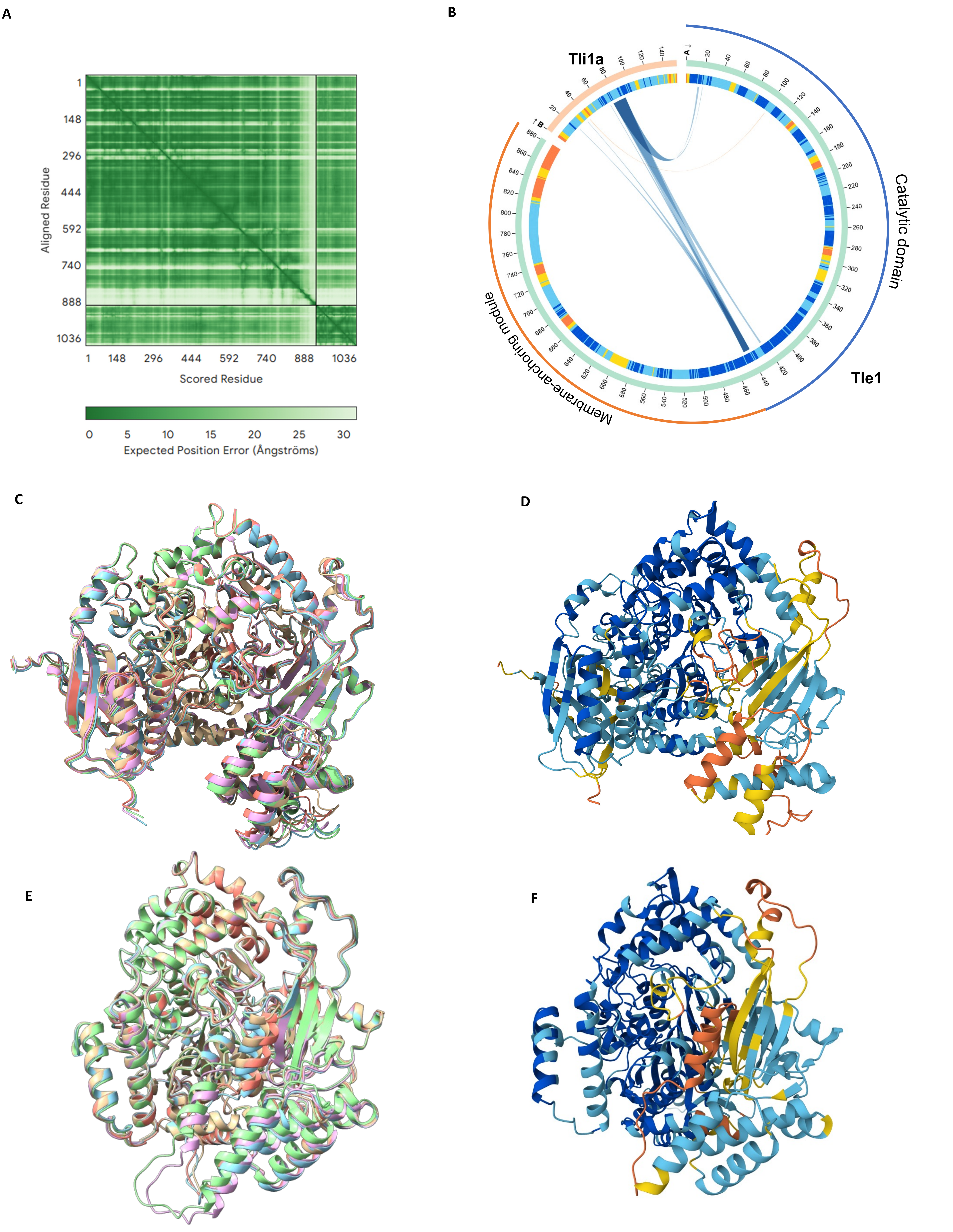


**Fig. S5 : Confidence metrics for the AlphaFold 3 models from (A-D) Fig. 3E (a monomer of Tle1 and a monomer of Tli1a) and (E-F) from Fig. 3F (a monomer of Tle1). (A)** AlphaFold3 predicted aligned error (PAE) plots. The color bar corresponds to expected position errors (Å). **(B)** AlphaBridge diagram of the Tle1-Tli1a complex. The outer and inner rings show the number of residues and the pLDDT for each chain, respectively. Colors in the outer rings correspond to pLDDT confidence scores: blue (very high), cyan (high), yellow (low), and orange (very low). **(C,E)** Overlay of the 5 best AlphaFold3 models. **(D,F)** AlphaFold3 predicted structure colored by prediction confidence (pLDDT) : blue (very high, pLDDT >90), cyan (high, 70> pLDDT> 90), yellow (low, 50> pLDDT> 70), and orange (very low, pLDDT <50).

**Supporting information Figure 5**

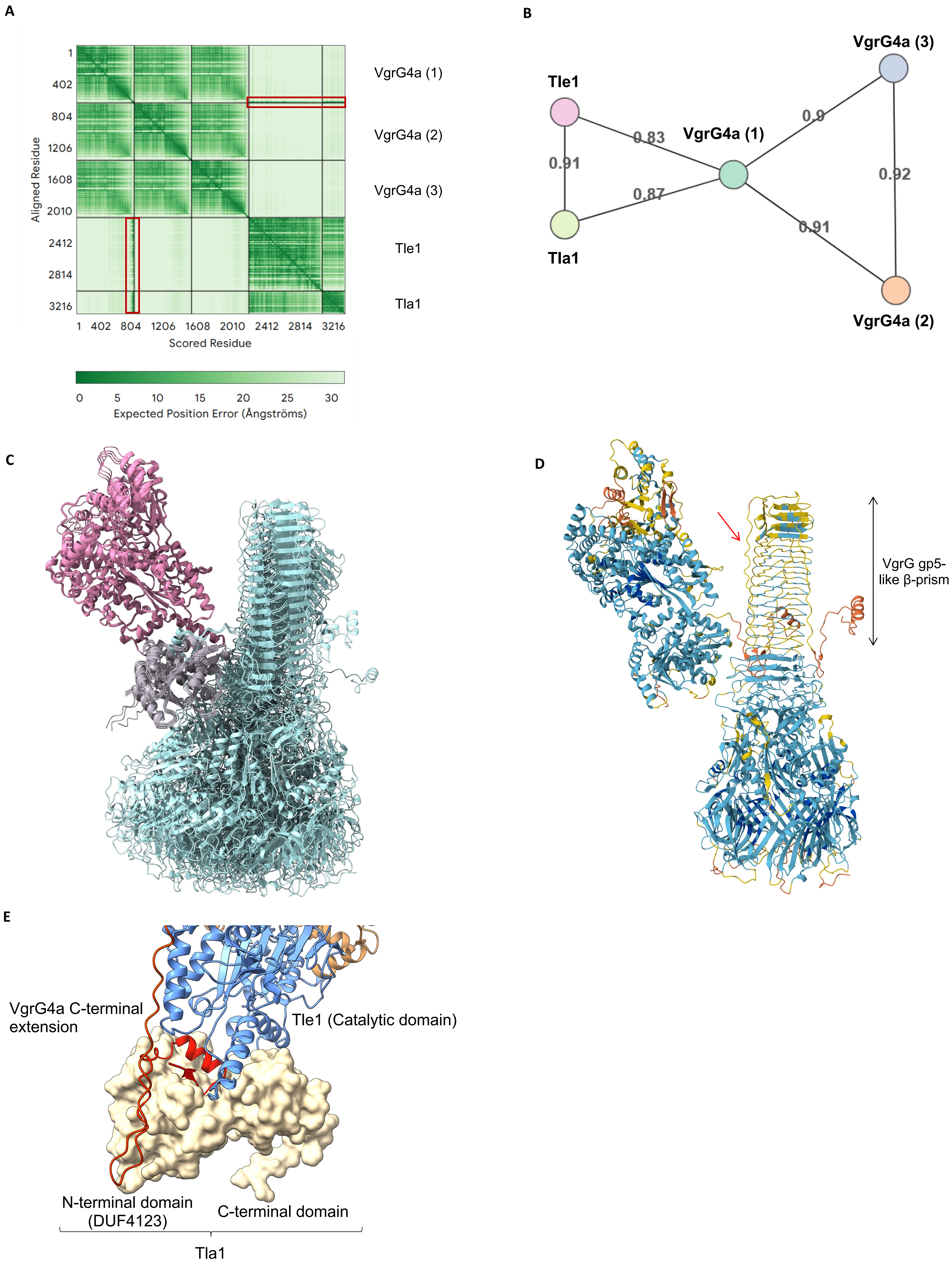


**Fig. S5 : Confidence metrics for the AlphaFold 3 model from Fig. 5 (trimer of VgrG4a, a monomer of Tle1 and a monomer of Tla1). (A)** AlphaFold3 predicted aligned error (PAE) plots. The color bar corresponds to expected position errors (Å). Red rectangles highlight the regions of high confidence (low error values) of the Tle1-VgrG4a (1) and Tla1-VgrG4a (1) interfaces. **(B)** Biomolecular network with the piCS (predicted interaction Confidence Score) indicated for each interfaces, along the edges of the model. **(C)** Overlay of the 5 best AlphaFold3 models. **(D)** AlphaFold3 predicted structure colored by prediction confidence (pLDDT) : blue (very high, pLDDT >90), cyan (high, 70> pLDDT> 90), yellow (low, 50> pLDDT> 70), and orange (very low, pLDDT <50). The red arrow highlights the flexible segment in the VgrG4a C-terminal extension, located near the gp5-like β-prism, which may adopt different positions. **(E)** Close-up view of the AlphaFold3 model from Fig. 4 showing Tla1 (surface representation, wheat), which is organized into two domains: the N-terminal DUF4123 domain connecting to the VgrG4a C-terminal extension (ribbons, red), and the C-terminal region connecting to the Tle1 catalytic domain (ribbons, blue).

**Supporting information Figure 6**

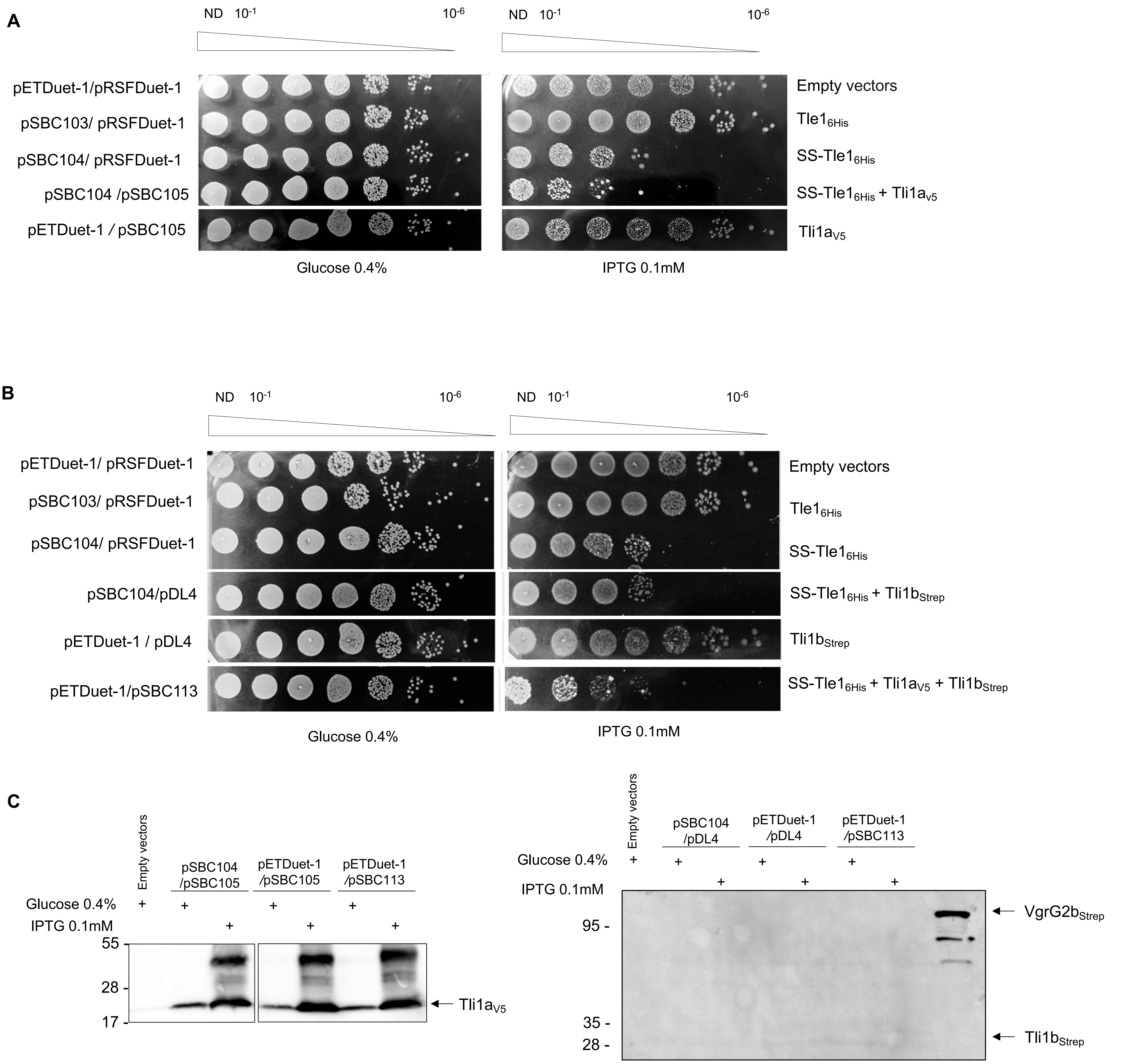


**Fig. S6 : Heterologous toxicity test in *E. coli.*** Cultures of BL21(DE3) pLysS producing Tle1$_{6His}$ (from pSBC103, line 2), SS-Tle1$_{6His}$(from pSBC104, line 3) with **(A)** Tli1a$_{V5}$ (from pSBC105) (lines 4,5), **(B)** Tli1b$_{Strep}$ (from pDL4) (lines 4,5) or Tli1a$_{V5}$ and Tli1b$_{Strep}$ (from pSBC113, expressed on the same transcript than *ss-tle1*) (line 6) were serially diluted (ND: not diluted to $10^{-6}$). Drops of each culture are deposited on LB agar glucose 0.4% medium to repress gene expression (left panel) or on LB agar IPTG 0.1 mM to induce gene expression (right panel). Strains containing the empty vectors pET22b(+) and pRSF-Duet-1 are used as controls (line 1). **(C)** Cells from panel (A and B) were collected and analyzed by Western-blot using an anti-V5 (Tli1a) or anti-Strep antibodies (Tli1b). VgrG2b$_{Strep}$ was used as a control for the anti-Strep antibodies.

**Supporting information Figure 7**

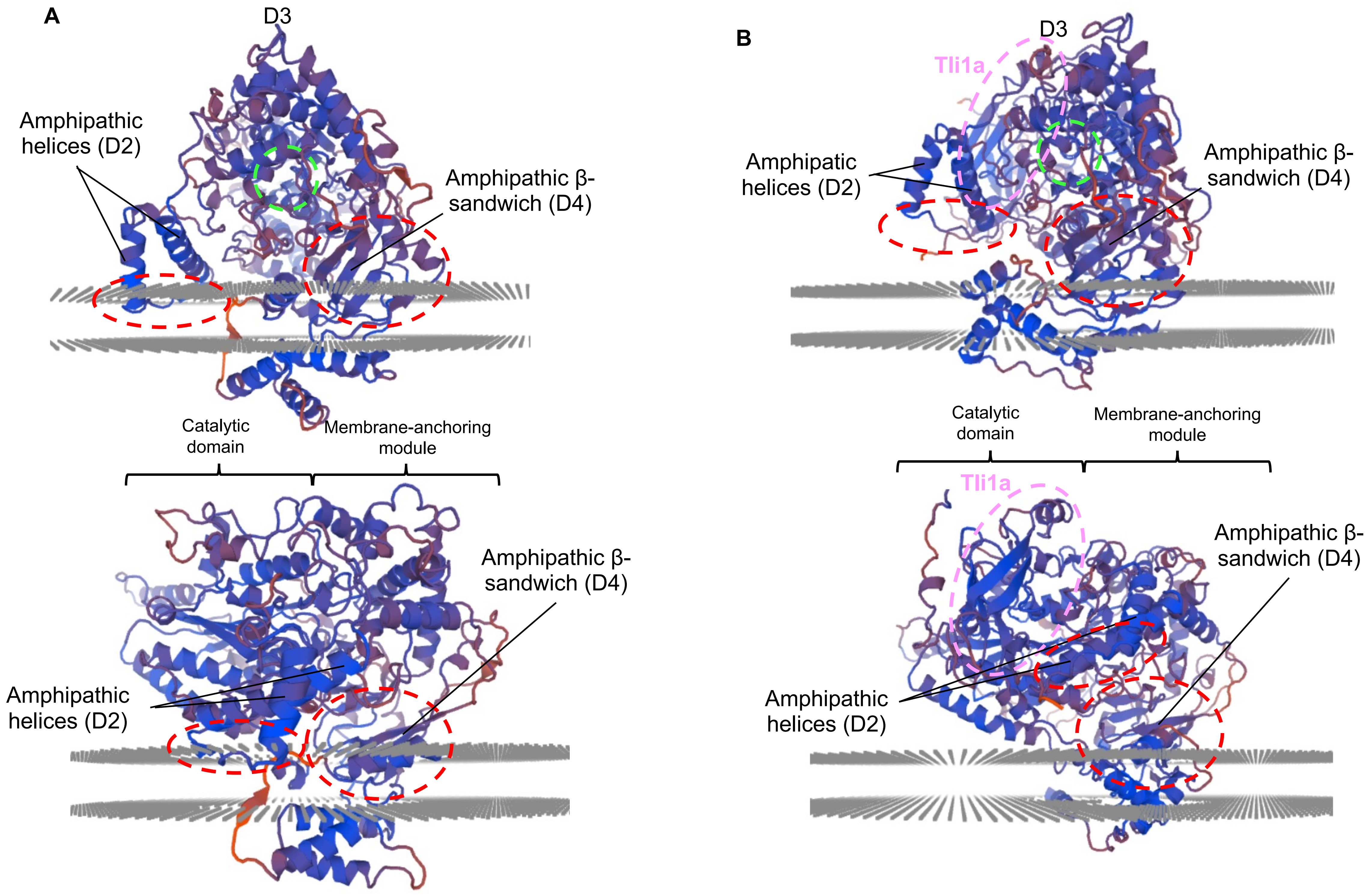


**Fig. S7 : Membrane insertion model of Tle1 in the presence of Tli1a.** The AlphaFold 3 Tle1 model (A) and the Tle1-Tli1a complex (B) were analyzed using the Swiss-Model QMEANBrane tool (Studer *et al.,* 2014). Top panels show the membrane-anchoring module, and bottom panels show a view of both Tle1 domains. The active site is indicated by a green dashed circle, and the loop between the two amphipathic helices of D2 and the amphipathic β-sandwich of D4, predicted to insert into the membrane, are indicated by a red dashed circle. Local QMEANBrane quality scores for membrane, interface, and soluble regions are shown (red : poor quality ; blue = high quality).

**Supporting information Figure 8**

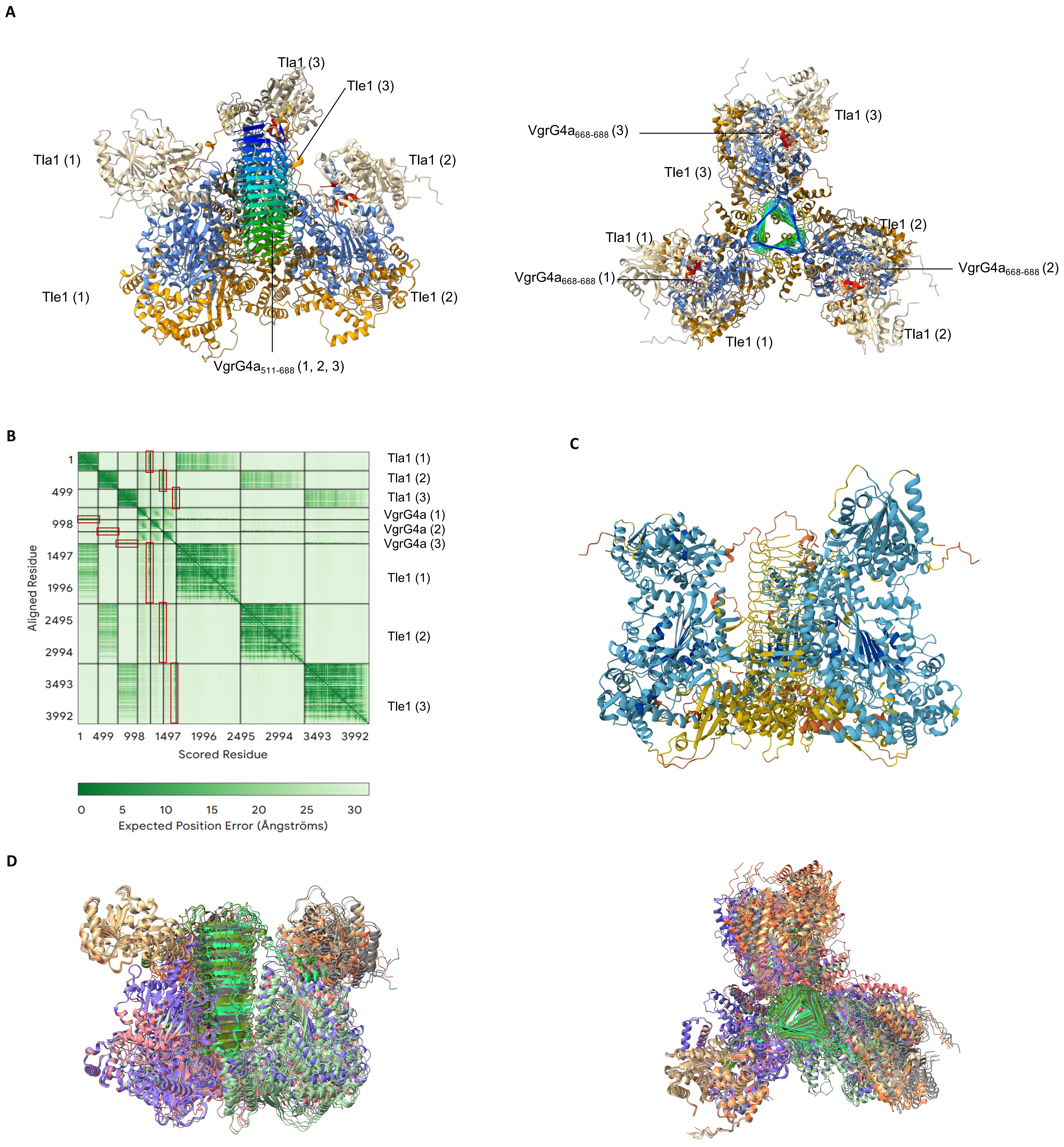


**Fig. S8: Each C-terminal extension of VgrG4a recruits a Tle1 and its chaperone Tla1 in a VgrG$_3$ (residues 511-688) –Tle1$_3$–Tla1$_3$ complex. (A)** AlphaFold3 predicted structure of the complex (left, front view and right, top view) between a trimer of VgrG4a (VgrG4a (1,2,3) in rainbow), three monomers of Tle1 (Tle1 (1,2,3), blue : catalytic domain, orange : membrane-anchoring module) and three monomers of Tla1 (Tla1 (1,2,3, wheat). **(B)** AlphaFold3 predicted aligned error (PAE) plots for Tle1$_3$- Tla1$_3$-VgrG4a$_3$ complex. The color bar corresponds to expected position errors (Å). The red rectangles highlight the confidence (low error values) of the Tle1$_3$-VgrG4a$_3$ (residues 511-688) and Tla1$_3$-VgrG4a$_3$ (residues 511-688) predicted interfaces. **(C)** AlphaFold3 predicted structure colored by prediction confidence (pLDDT) : blue (very high, pLDDT >90), cyan (high, 70> pLDDT> 90), yellow (low, 50> pLDDT> 70), and orange (very low, pLDDT <50). **(D)** Overlay of the 5 best AlphaFold3 models.
